# A Single Membrane Protein Required for Atrial Secretory Granule Formation

**DOI:** 10.1101/2020.03.08.982777

**Authors:** Nils Bäck, Raj Luxmi, Kathryn G. Powers, Richard E. Mains, Betty A. Eipper

## Abstract

The discovery of atrial secretory granules and the natriuretic peptides stored in them identified the atrium as an endocrine organ. Although neither atrial nor brain natriuretic peptide (ANP, BNP) is amidated, the major membrane protein in atrial granules is Peptidylglycine α-Amidating Monooxygenase (PAM), an enzyme essential for amidated peptide biosynthesis. Mice lacking cardiomyocyte PAM (*Pam*^Myh6-cKO/cKO^) are viable, but a gene dosage-dependent drop in atrial ANP and BNP content occurred. Ultrastructural analysis of adult *Pam*^Myh6-cKO/cKO^ atria revealed a 20-fold drop in the volume fraction of secretory granules and a decrease in peripherally localized Golgi complexes. When primary cultures of *Pam*^*0-Cre*-cKO/cKO^ atrial myocytes (PAM floxed, no Cre recombinase) were transduced with Cre-GFP lentivirus, PAM protein levels dropped, followed by a decline in proANP levels. Expression of exogenous PAM in *Pam*^Myh6-cKO/cKO^ atrial myocytes produced a dose-dependent increase in proANP content. Strikingly, rescue of proANP content did not require the monooxygenase activity of PAM. Unlike many prohormones, atrial proANP is stored intact and its basal secretion is stimulated by drugs that inhibit Golgi-localized Arf activators. Increased basal secretion of proANP was a major contributor to its reduced levels in *Pam*^Myh6-cKO/cKO^ myocytes; the inability of these drugs to inhibit basal proANP secretion by *Pam*^Myh6-cKO/cKO^ myocytes revealed a role for COPI-mediated recycling of PAM to the endoplasmic reticulum. Analysis of atrial coated vesicles and the ability PAM to make fluorescently-tagged proANP accumulate in the *cis*-Golgi region of cells lacking secretory granules revealed a non-catalytic role for PAM in soluble cargo trafficking early in the secretory pathway.

**Significance:** Transmission electron microscopy of atrial cardiomyocytes revealed dense granules resembling those in endocrine cells and neurons, leading to the discovery of the natriuretic peptides stored in these granules. Subsequent studies revealed features unique to atrial granules, including high level expression of Peptidylglycine α-Amidating Monooxygenase (PAM), an enzyme required for the synthesis of many neuropeptides, but not for the synthesis of natriuretic peptides. The discovery that atrial myocytes lacking PAM are unable to produce granules and that PAM lacking its monooxygenase activity can rescue granule formation provides new information about the proANP secretory pathway. A better understanding of the unique features of atrial cell biology should provide insight into atrial fibrillation, the most common cardiac arrhythmia, atrial amyloidosis and heart failure.

## INTRODUCTION

In response to stretch, sympathetic input and hormones like endothelin-1, atrial myocytes release natriuretic peptides essential for fluid volume homeostasis (1, 2). The pro-atrial natriuretic peptide (proANP) containing granules in atrial myocytes, which look much like the granules in insulin-producing β-cells and proopiomelanocortin (POMC)-producing corticotropes, exhibit important differences. Atrial granules accumulate near the Golgi complex and store their major product, proANP, intact. The subtilisin-like endoproteases responsible for producing product peptides from inactive precursors in neurons and endocrine cells are not expressed in cardiomyocytes (3). Instead, Corin cleaves proANP at the time of secretion (1, 2, 4). While calcium influx or intracellular release plays a major role in the stimulated secretion of islet, pituitary and chromaffin cell hormones, it is not essential for atrial proANP secretion (5, 6). Specific aspects of the secretion of many peptide hormones (7-9) are inhibited by Brefeldin A (BFA), a fungal metabolite which inhibits the Golgi localized GDP/GTP exchange factors (GEFs) that activate Arf family GTP binding proteins (10-12). In contrast, atrial myocyte secretion of ANP is stimulated by BFA (13, 14).

Natriuretic peptides, proANP and pro-brain natriuretic peptide (proBNP), are the major atrial granule content proteins. Granule membrane proteins play essential roles in granule formation and maturation, cargo protein post-translational processing, luminal pH maintenance, ion transport and cytoskeletal interactions. Strikingly, peptidylglycine α-amidating monooxygenase (PAM), a Type I transmembrane enzyme that converts peptidylglycine substrates (luminal peptides with a COOH-terminal Gly) into amidated products, is the major atrial granule membrane protein (15, 16). For many neuroendocrine peptides, amidation is essential to function (17). However, neither ANP nor BNP is amidated, suggesting that PAM performs a non-catalytic, structural role in atrial granules (15, 16). In support of this idea, PAM has been reported to interact with proANP in a manner dependent on the proANP calcium-binding site (18).

Genetic studies have linked both *NPPA* and *PAM* to heart disease. A mutation in *NPPA* was identified as a causative factor in familial, early onset atrial fibrillation (19). In isolated atrial amyloidosis, which is associated with congestive heart failure and atrial fibrillation, ANP and its N-terminal fragment are major amyloid fibril components (20). PAM has been identified as a risk factor for type 2 diabetes (21, 22), coronary heart disease (21, 23) and hypertension (24). Although atrial fibrillation, the most common arrhythmia in the elderly and a leading cause of stroke, is accompanied by changes in the organization of the Golgi complex and microtubule network in atrial myocytes (25-27), our limited understanding of the unique features of these cells limits our ability to manipulate them in a therapeutically useful manner.

Mice unable to express PAM in cardiomyocytes were generated by crossing mice expressing a floxed allele of *Pam* (*Pam*^*0-Cre*-cKO/cKO^) to mice expressing Cre recombinase under control of the myosin heavy chain 6 (*Myh6*) promoter (3, 28). *Pam*^Myh6-cKO/cKO^ mice have dramatically reduced levels of proANP in their atria and mature ANP in their sera (3). These decreases cannot be attributed to a decline in *Nppa* mRNA levels. We used ultrastructural, biochemical and pharmacological approaches to demonstrate the inability of *Pam*^Myh6-cKO/cKO^ atrial myocytes to store selected soluble cargo proteins in granules. Our exploration of the underlying mechanism highlighted unique features of the secretory pathway in atrial myocytes, revealing a role for PAM in prolonging the half-life of proANP by protecting it from basal secretion and degradation.

## RESULTS

### Secretory granules are rarely seen in the atria of *Pam*^Myh6-cKO/cKO^ mice

We turned to transmission electron microscopy to determine whether the drop in proANP levels observed in the atria of *Pam*^Myh6-cKO/cKO^ mice (3) were accompanied by a decrease in the number of secretory granules. The cytoplasmic volume of atrial myocytes is largely filled by myofilaments and mitochondria (29). To obtain representative images of the organelles of interest, randomly oriented tissue blocks from the left atria of three wildtype and three *Pam*^Myh6-cKO/cKO^ mice were examined (**Methods**). Myofilaments and mitochondria did not differ. Perinuclear Golgi complexes (G) and nearby secretory granules (arrowheads) were readily observed in control myocytes (**Fig. 1A**). Although perinuclear Golgi complexes were present in *Pam*^Myh6-cKO/cKO^ atria, secretory granules were rarely observed in their vicinity (**Fig. 1B**). Irregularly shaped granules (inset to **Fig. 1B**) were more common in the perinuclear Golgi region of *Pam*^Myh6-cKO/cKO^ atria than in control atria (52.7 ± 4.0% of the granules in *Pam*^Myh6-cKO/cKO^ atria vs. 11.7 ± 0.7% in control atria; ± SE, p < 0.001).

**Fig. 1.**
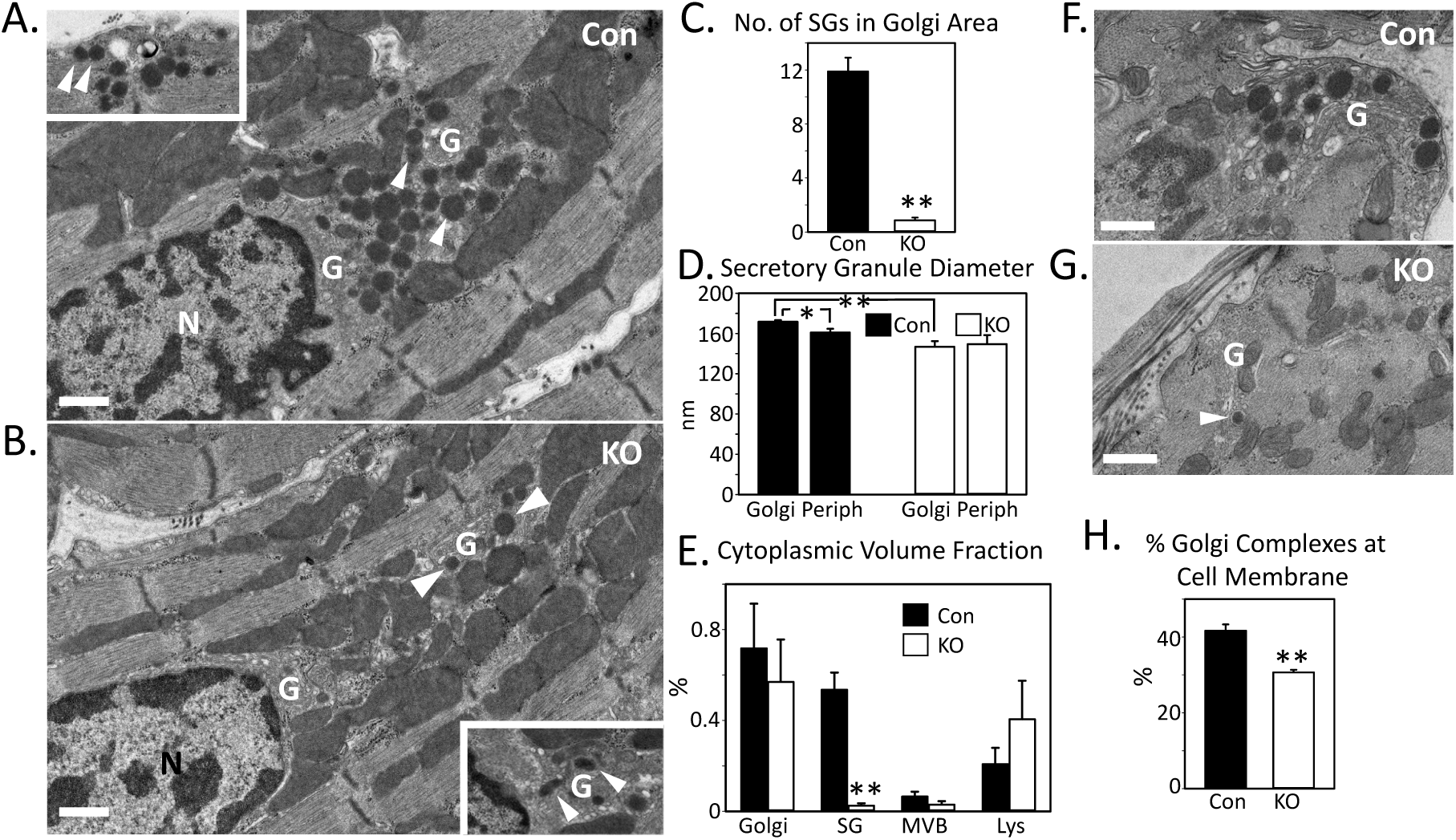
Secretory granules are rarely seen in the atria of *Pam*^Myh6-cKO/cKO^ mice. **A.** Images of control (**A**) and *Pam*^Myh6-cko/cko^ (**B**) atria including nucleus (N) and Golgi (G) complex; secretory granules (SGs) are marked with *arrowheads. Inset* in (**A**) shows SGs at the cell periphery. Inset in (**B**) shows Golgi membranes and irregular SGs. Quantification of SG number in sections including the Golgi complex (Golgi membranes bordered by mitochondria and myofilament bundles) (**C**), SG diameter in sections including the Golgi or peripheral membranes (**D**) and cytoplasmic volume fractions for Golgi, SGs, multivesicular bodies (MVB) and lysosomes (Lys) (**E**) in control and *Pam*^Myh6-cKO/cKO^ (KO) atria. In **C, D** and **E**, ** p < 0.001 and * p < 0.05. In **H**, ** p < 0.005. **F, G.** Images of Golgi complexes at the cell membrane in control and *Pam*^Myh6-cko/cko^ atria. *Arrowhead* in **G** shows a SG. **H.** Quantification of the percentage of Golgi complexes at the cell membrane. Scale bars, 500 nm.

Quantification of the number of secretory granules observed in transections of the Golgi complex revealed a 13-fold decrease in the left atria of *Pam*^Myh6-cKO/cKO^ mice (**Fig. 1C**); the drop in granule number exceeded the 3-fold drop in proANP content observed in *Pam*^Myh6-cKO/cKO^ atria. In control atrial myocytes, granules located in the perinuclear Golgi region had a larger diameter than granules located near the sarcolemma. The granules in the perinuclear Golgi region in *Pam*^Myh6-cKO/cKO^ myocytes had a smaller diameter than similarly localized granules in control cells, but their diameters did not differ from peripherally localized granules (**Fig. 1D**).

Atrial myocyte cell size was unaltered in *Pam*^Myh6-cKO/cKO^ mice; the width of cell transections averaged 5.7 ± 0.1 µm in control cells and 6.1 ± 0.1 µm in *Pam*^Myh6-cKO/cKO^ cells; the nuclear volume fraction was also unaltered (3.6 ± 0.5% in control cells; 2.8 ± 0.5% in *Pam*^Myh6-cKO/cKO^ cells). Morphometric analysis of the cytoplasmic volume occupied by Golgi complex, secretory granules, multivesicular bodies (MVBs) and lysosomes revealed a 20-fold decrease for secretory granules in *Pam*^Myh6-cKO/cKO^ cardiomyocytes (**Fig. 1E**). None of these organelles occupied more than 1% of the cytoplasmic volume.

In addition to prominent perinuclear Golgi complexes and granules, Golgi membranes were observed near the periphery of control and *Pam*^Myh6-cKO/cKO^ myocytes (**Fig. 1F, G**) (25). In control cells, 40% of the Golgi membranes were located near the cell surface; a smaller percentage of the Golgi complexes were located near the cell membrane in *Pam*^Myh6-cKO/cKO^ myocytes (**Fig. 1H**). As for the perinuclear Golgi complexes, secretory granules were prevalent near the peripheral Golgi complexes in control cells but were rarely observed in this region in *Pam*^Myh6-cKO/cKO^ myocytes.

### Endosomal/lysosomal organelles in the atria of *Pam*^Myh6-cKO/cKO^ mice

In pituitary cells, PAM that reaches the plasma membrane is retrieved via clathrin-mediated endocytosis and is either recycled to secretory granules or degraded (30, 31). The endocytic trafficking of PAM is regulated and involves its entry into the intraluminal vesicles of multivesicular bodies (MVB). For this reason, we examined MVBs in control and *Pam*^Myh6-cko/cko^ atrial myocytes. In both genotypes, MVBs were observed near the Golgi complex and secretory granules (SG) (**Fig. 2A-D**, arrowheads). Irregularly shaped, dark staining lysosomes (Lys) were seen close to the Golgi complex in both control and *Pam*^Myh6-cKO/cKO^ atrial myocytes (**Fig. 2E, F**). Residual bodies, which release membranous aggregates, did not vary with genotype (**Fig. 2G**, arrowheads). In both control and *Pam*^Myh6**-**cKO/cKO^ myocytes, MVBs containing electron dense material resembling the cores of secretory granules were occasionally observed (**Fig. 2H, I**), consistent with the occurrence of crinophagy (32).

**Fig. 2.**
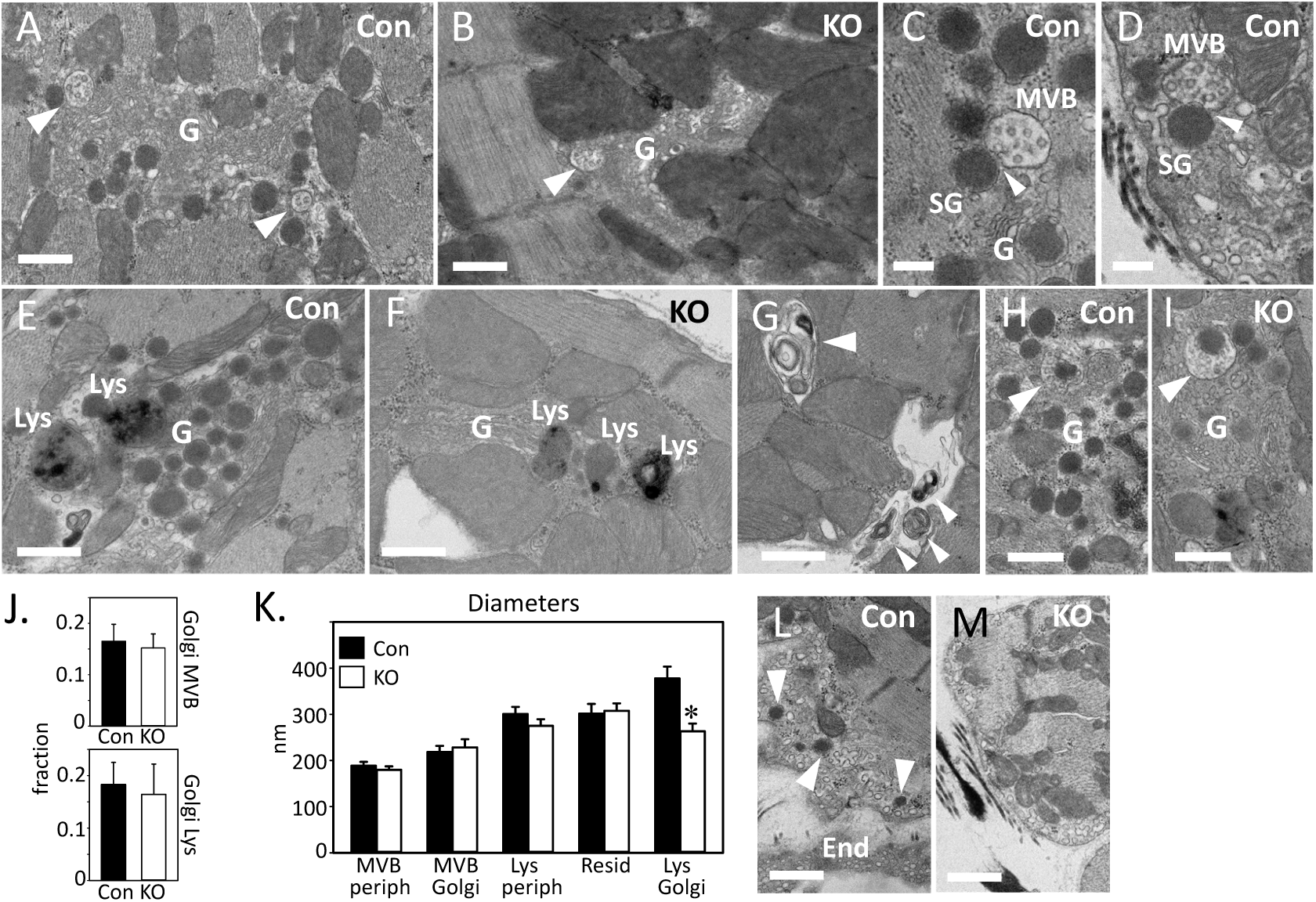
MVBs and lysosomes in control and *Pam*^Myh6-cKO/cKO^ atria. MVBs (*arrowheads)* in the Golgi (G) region in control **(A, C, D)** and *Pam*^Myh6-cKO/cKO^ **(B)** atrial myocytes. In controls, MVBs interacted closely with SGs in the Golgi area (**A, C**) and in the periphery (**D**). Dark, amorphous lysosomes (Lys) were seen near the Golgi complex in control (**E**) and *Pam*^Myh6-cKO/cKO^ (**F**) atrial myocytes. Residual bodies in a control myocyte are shown (**G**, *small arrowheads*). In both control (**H**) and *Pam*^Myh6**-**cKO/cKO^ (**I**) atrial myocytes, SG destruction by crinophagy occurs (*arrowheads*). The number of MVBs and Lys (**J)** located in the Golgi region did not vary with genotype. (**K**) MVB, Lys and residual body (Resid) diameters were measured for organelles located peripherally (periph) or near the Golgi complex; Golgi-localized Lys were smaller in *Pam*^Myh6-cKO/cKO^ atrial myocytes than in control cells (**\*** p < 0.005). The peripheral rim of cytoplasm was occupied by an extensive array of vesicles, vacuoles and invaginations in control and KO cells **(L**,**M)**; in control cells, SGs (arrowheads) were regularly observed. End, endothelial cell. Scale bars, **A, B, E-G, L-M**, 500 nm; **C, D**, 200 nm.

The fraction of Golgi complex transections containing MVBs or lysosomes did not differ in control and *Pam*^Myh6-cKO/cKO^ myocytes (**Fig. 2J, K**). The diameters of peripheral and Golgi localized MVBs, peripheral lysosomes and residual bodies did not differ based on genotype, but Golgi localized lysosomes were smaller in *Pam*^Myh6-cKO/cKO^ myocytes (**Fig. 2K**); the observed decrease in diameter would account for a 3-fold drop in the volume of Golgi-localized lysosomes. The peripheral rim of cytoplasm was filled with an extensive array of invaginations, vesicles and tubules (**Fig. 2L, M**). Ultrastructurally, it was not possible to distinguish T-tubules, sarcoplasmic reticulum, caveolae and endocytic vesicles. The volume fraction of these structures was identical in control and *Pam*^Myh6-cKO/cKO^ cardiomyocytes (0.96 ± 0.19% and 0.95 ± 0.29%, respectively). Based on an unbiased transmission electron microscopic analysis of secretory pathway organelles in atrial myocytes from control and *Pam*^Myh6-cKO/cKO^ mice, the major difference was the almost total absence of atrial granules in myocytes lacking PAM.

### Granule cargo protein levels are reduced in *Pam*^Myh6-cKO/+^ atria and further reduced in *Pam*^Myh6-cKO/cKO^ atria

Eliminating the expression of *Pam* in atrial cardiomyocytes resulted in decreased levels of proANP in the atrium (3). To determine the effect of loss of a single copy of *Pam*, we quantified PAM and proANP protein in atria from control mice, heterozygous mice (*Pam*^Myh6-cKO/+^) and *Pam*^Myh6-cKO/cKO^ mice (**Fig. 3A**). As observed in adult *Pam*^+/-^ mice, levels of intact PAM protein in the atria of *Pam*^Myh6-cKO/+^ mice were less than half of control values (33). PAM protein was undetectable in the atria of *Pam*^Myh6-cKO/cKO^ mice. Levels of proANP fell below control values in atria from *Pam*^Myh6-cKO/+^ mice and fell further in atria from *Pam*^Myh6-cKO/cKO^ mice (**Fig. 3A**).

**Fig. 3.**
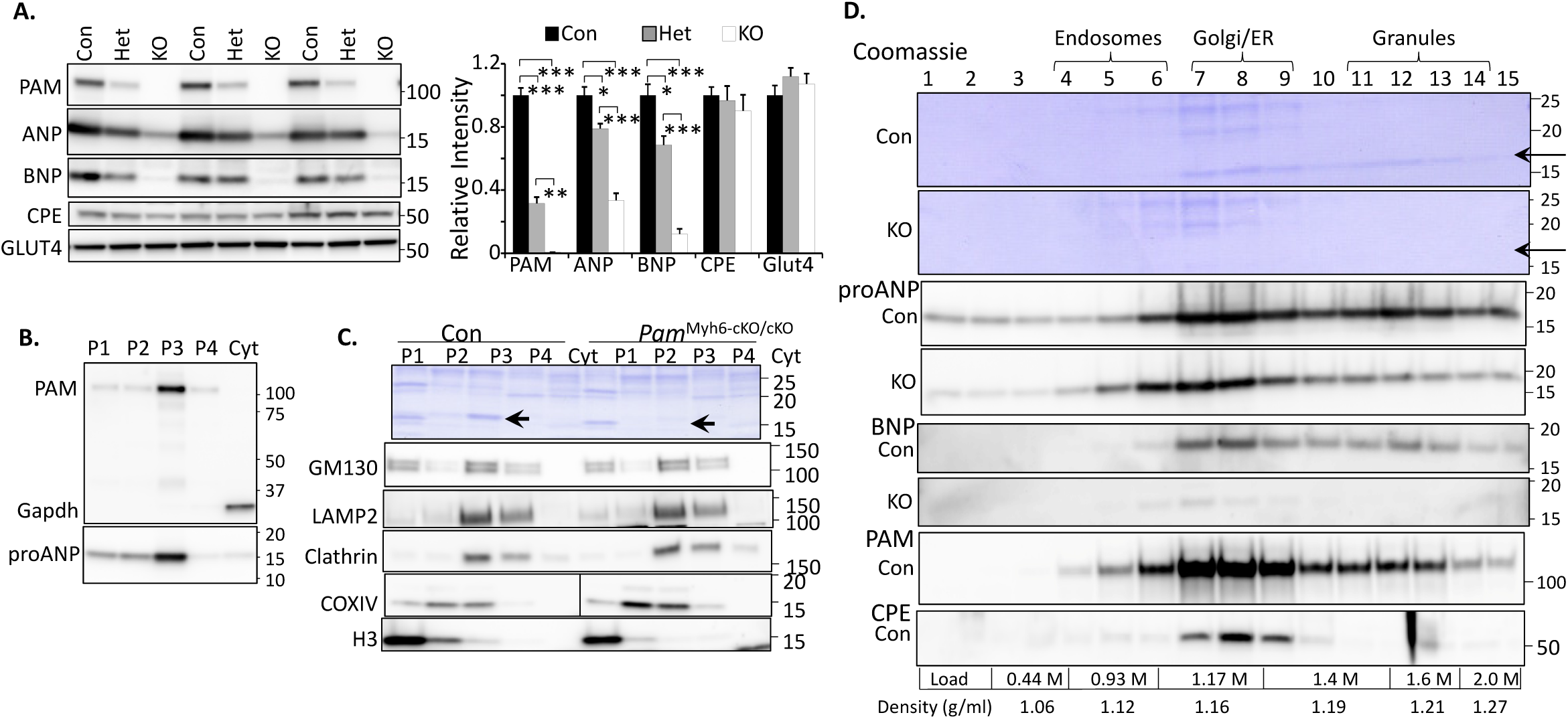
PAM exerts a dosage-dependent, cargo-specific effect on atrial granules. **A.** Atrial lysates (20 μg) prepared from control (Con), heterozygous (Het, *Pam* ^Myh6-cKO/+^) and homozygous (KO, *Pam*^Myh6-cKO/cKO^) adult male and female mice were fractionated by SDS-PAGE. Expression of PAM, three soluble secretory pathway cargo proteins (ANP, BNP and CPE) and one membrane protein (GLUT4) was evaluated; *, p<0.01; **, p<0.001, ***, p<0.0001. **B.** Western blot analysis was used to determine the localization of PAM, Gapdh and proANP in subcellular fractions prepared from control adult male atria by differential centrifugation. **C.** Subcellular fractions (equal amounts of protein) from control and *Pam*^Myh6-cKO/cKO^ atria were fractionated by SDS-PAGE. The bottom part of the Coomassie stained PVDF membrane is shown, with arrows marking the expected migration position of proANP. Western blot analysis identified fractions enriched in Golgi membranes (GM130), lysosomes (LAMP2), trafficking vesicles (clathrin), mitochondria (COX IV) and nuclei (Histone H3, H3). **D.** Sucrose density gradient fractionation of P3 fractions from control and *Pam*^Myh6-cKO/cKO^ (KO) atria. The bottom part of the Coomassie-stained PVDF membranes is shown, with arrows marking the expected migration position of proANP. The molarities and densities of the sucrose solutions used to form the step gradient appear at the bottom. Western blot analyses for proANP and BNP from Con and KO atria are shown. PAM was detected in the ER/Golgi and granule-rich regions of the control gradient; only 20.6 ± 0.7% (n = 3) of the PAM protein on the P3 gradient was recovered from the granule-rich region, indicating that proANP was more effectively packaged in granules than PAM (p < 0.001). Similar results were obtained in two additional direct comparisons of WT and *Pam*^Myh6-cKO/cKO^ P3 fractions and in multiple analyses of WT P3 samples.

In addition to proANP, atrial granules contain proBNP and carboxypeptidase E (CPE) (15). As observed for proANP, levels of proBNP protein fell in *Pam*^Myh6-cKO/+^ atria and fell further in *Pam*^Myh6-cKO/cKO^ atria (**Fig. 3A**). In contrast, levels of CPE, another soluble cargo protein, were unaltered in atria lacking *Pam*. Glut4 (Slc2a4), a glucose transporter that is rapidly moved to the sarcolemma in response to insulin and enters proANP-containing granules after endocytic removal (34) was also examined. Levels of Glut4 protein were unaltered in atria lacking *Pam* (**Fig. 3A**). To determine the specificity of the changes observed, expression of markers for different organelles was also evaluated; no changes occurred (**Fig. S1A**). Levels of several of the cytosolic and membrane proteins previously identified as PAM interactors were also unchanged in atria lacking *Pam*, but Vamp4 levels were reduced (**Fig. S1B**) (35, 36). The lack of PAM protein exerted a cargo-specific effect on soluble secretory pathway proteins identified in atrial granules.

### Subcellular fractionation of mouse atrium

Secretory granule volume fell 20-fold in the atria of *Pam*^*Myh6-cKO/cKO*^ mice while atrial proANP levels fell only 3-fold. To determine the fate of granule content proteins in *Pam*^Myh6-cKO/cKO^ atria, existing methods were modified to allow simultaneous evaluation of multiple organelles (16, 37, 38). Homogenization and differential centrifugation conditions were optimized for recovery of PAM and proANP in particulate fractions, with Gapdh, but very little proANP, in the cytosol (**Fig. 3B**). Intact PAM and proANP were enriched in the P3 fraction, with lower levels of intact PAM in P4, the fraction expected to contain endosomes. We next compared subcellular fractions prepared from the atria of control and *Pam*^Myh6-cKO/cKO^ mice (**Fig. 3C**). Coomassie staining revealed a prominent 16 kDa band, the mass of proANP and proBNP, in the P3 fraction of control, but not *Pam*^Myh6-cKO/cKO^, mice (arrows in upper panel). Nuclear, mitochondrial, lysosomal, Golgi and vesicular trafficking markers were similarly localized in both genotypes (**Fig. 3C**).

Sucrose density gradient centrifugation was used to separate atrial granules from lighter membranous organelles (39) (**Fig. 3D**). Antisera to ER, Golgi and endosomal markers revealed similar profiles for P3 pellets from control and *Pam*^Myh6-cko/cko^ atria (**Fig. S2**), with no clear separation of ER and Golgi markers. In P3 pellets from control atria, proANP was localized to the ER/Golgi region and to denser regions of the gradient, consistent with its presence in granules (**Fig. 3D**). The denser fractions accounted for 35.4 ± 2.5% of the total P3 proANP signal in control atria (n = 4), but only 25.7 ± 2.5% in *Pam*^Myh6-cKO/cKO^ atria (n = 2; p < 0.005). Coomassie staining revealed a prominent band at approximately 16 kDa only in the control P3 fraction (**Fig. 3D**, arrow); the granule region accounted for 33.9 ± 4.7 % (n = 3) of the total 16 kDa Coomassie signal. As for proANP, Western blot analysis revealed peaks of proBNP in both the ER/Golgi and granule-enriched regions of the P3 gradient (**Fig. 3D**). In P3 fractions prepared from *Pam*^Myh6-cKO/cKO^ atria, proBNP was detected in the ER/Golgi region, but not in the granule-containing region.

To evaluate the specificity of this effect, the subcellular localization of CPE was determined. In both control (**Fig. 3D**) and *Pam*^Myh6-cKO/cKO^ (not shown) atria, CPE was enriched in the ER/Golgi rich fractions, not in the dense fractions containing the natriuretic peptides. Based on both transmission EM and biochemical fractionation, atrial myocyte PAM is essential for normal proANP and proBNP storage. With an almost complete loss of morphologically identifiable granules in *Pam*^Myh6-cKO/cKO^ atria, proANP must occupy a different part of the secretory pathway.

### Expression of PAM in *Pam*^Myh6-cKO/cKO^ atrial myocytes produces a dose-dependent increase in proANP levels

Expression of PAM in rodent cardiomyocytes is readily detectable after embryonic day 14 (40). In order to determine whether the absence of atrial granules in adult *Pam*^Myh6-cKO/cKO^ mice reflected a developmental defect, we prepared primary cultures from the atria of mice expressing two copies of the floxed *Pam* allele but no Cre-recombinase (*Pam*^*0-Cre*-cko/cko^), meaning that PAM protein levels were normal. These atrial cultures were transduced with lentiviruses expressing GFP (control) or Cre-GFP (**Fig. 4A, B**). After Cre-GFP-mediated excision of *Pam* exons 2+3, the decline in PAM protein reflects both the half-life of its mRNA and the half-life of the PAM protein. Levels of PAM, proANP and Gapdh were unaltered in cultures expressing virally-encoded GFP. In cultures expressing Cre-GFP, levels of PAM declined by more than a factor of 2 within 4 days and by more than a factor of 10 within 6 days. As expected, levels of proANP fell more slowly, dropping about 4-fold in 9 days (**Fig. 4B**). The time course over which proANP levels declined indicated that the ability of atrial myocytes to store proANP was dependent on the continued presence of PAM.

**Fig. 4.**
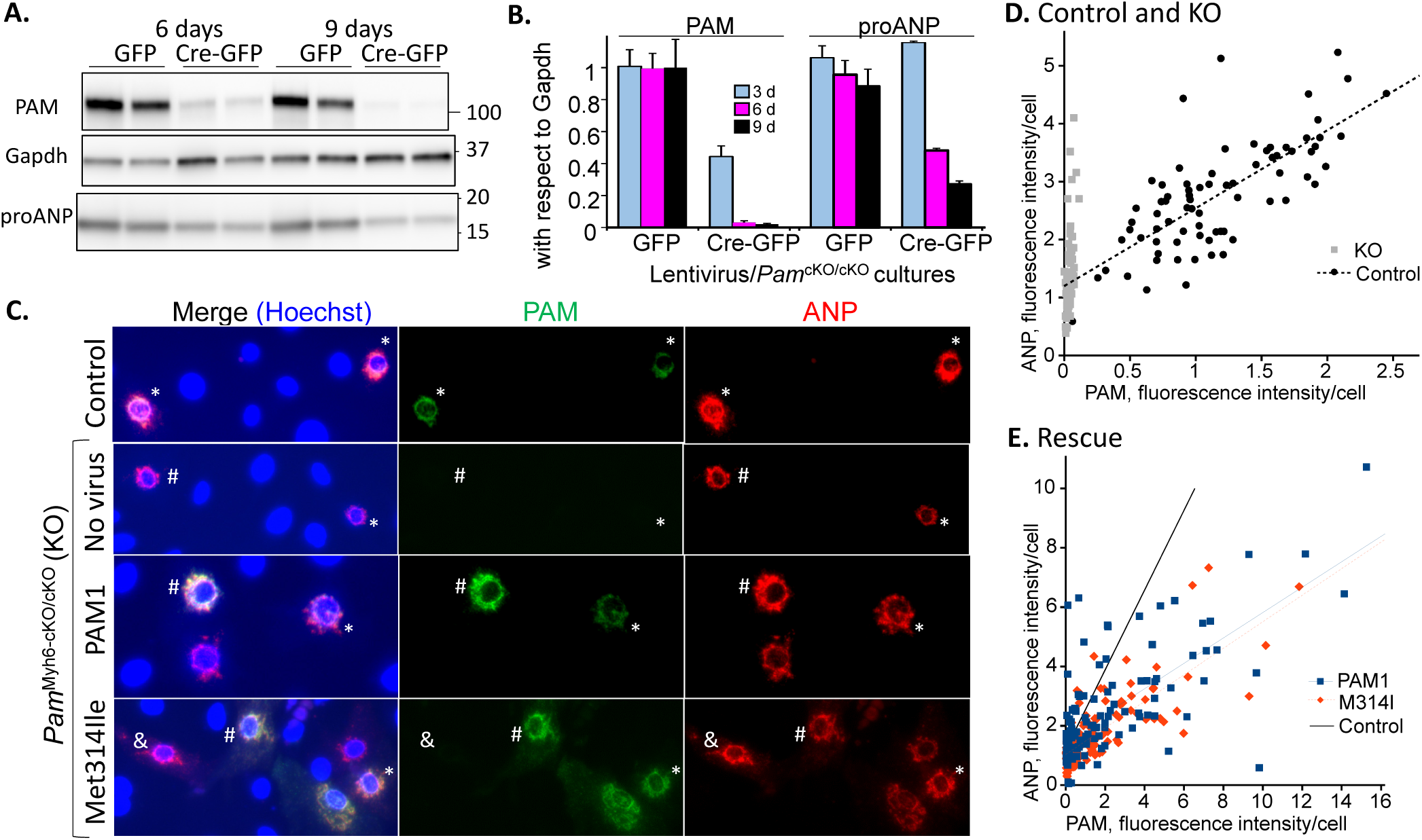
Expression of PAM controls the ability of *Pam*^Myh6-cko/cko^ myocytes to store proANP. **A.** Atrial cultures prepared from *Pam*^cKO/cKO^ pups were spinoculated with virus encoding GFP or Cre-GFP. After 3 (not shown), 6 or 9 days, cultures were harvested and equal amounts of cell protein (2.5 μg) were fractionated by SDS-PAGE; Western blot analysis for PAM (JH629), Gapdh and proANP revealed loss of PAM and proANP in cultures spinoculated with the Cre-GFP virus. **B.** Immunoblot signals were densitized, with PAM and proANP levels normalized to Gapdh; data were normalized to levels on the first harvest day. The experiment was repeated with similar results. **C.** Cultures were prepared at the same time from control and *Pam*^Myh6-cKO/cKO^ (KO) pups; control cultures were not spinoculated. *Pam*^Myh6-cKO/cKO^ cultures were analyzed without spinoculation (None) or after spinoculation with lentiviruses encoding PAM1 or PAM1/Met314Ile. Seven days later, cultures were fixed and stained simultaneously for ANP (goat ANP antibody; Alexa633 secondary, shown in red), PAM (rabbit PAM exon A antibody; Alexa488 secondary, shown in green) and nuclei (Hoechst stain; blue). Cultures were photographed under identical conditions; the exposure time for each fluorophore was adjusted to avoid saturation. Myocytes expressing higher and lower levels of PAM are indicated by an # and an *, respectively; myocytes with ANP but no PAM are marked by a &. **D. Control and KO.** Integrated total intensities for PAM and ANP staining were determined using FuJi and a locally determined background measurement for each cell. A range of PAM and ANP staining intensities was observed in control cells, clustered around a best-fit linear relationship (black circles). Variable ANP staining intensities were also observed in *Pam*^Myh6-cKO/cKO^ cells, but no PAM staining was detected (grey squares). X and Y axes are arbitrary linear scales for reporting integrated total intensity per cell. The average integrated ANP staining intensity/cell in control atrial myocytes was cut in half in myocytes lacking PAM (2.77 ± 0.11 vs. 1.40 ± 0.08). **E. Rescue.** Spinoculated cultures of *Pam*^Myh6-cKO/cKO^ cells expressing PAM1 showed an increase in ANP staining intensity as PAM staining intensity increased (blue squares). A similar increase in ANP staining intensity was observed in *Pam*^Myh6-cko/cko^ cells expressing PAM1/Met314Ile (M314I, orange diamonds).

We next wanted to determine whether expression of exogenous PAM in atrial myocytes lacking any PAM could rescue proANP levels to those in control myocytes. The high levels of PAM expression, despite the absence of high levels of any amidated peptide, suggested that PAM might play a non-catalytic role in the atrium. To test this hypothesis, lentiviruses encoding rat PAM1 and rat PAM1/Met314Ile, a catalytically inactive mutant in which the monooxygenase active site methionine residue was replaced by isoleucine, were constructed (41). By expressing lentiviral encoded PAM1 and PAM1/Met314Ile in HEK293 cells, we verified the lack of PHM activity and normal levels of PAL activity in cells expressing PAM1/Met314Ile (**Fig. S3**).

To evaluate the effect of expressing exogenous PAM1 or PAM1/Met314Ile on single myocytes, cultures were fixed and stained for proANP and for PAM. To validate our methodology, we first compared the properties of atrial myocytes prepared from control (*Pam*^0-Cre-cKO/cKO^) and *Pam*^Myh6-cKO/cKO^ (KO) mice (**Fig. 4C**); based on the developmental stage at which the Myh6 promoter becomes active and the time at which PAM is first expressed in the rodent atrium, PAM expression should never be initiated in *Pam*^Myh6-cKO/cKO^ atrial myocytes (42). Using images captured under identical conditions, individual myocytes were evaluated for their content of PAM and proANP. In control myocytes, levels of both proANP and PAM in individual myocytes varied widely (**Fig. 4D**, black circles), as observed for several endocrine cell markers in single pancreatic β-cells (43). Quantification of both signals revealed a positive correlation (R^2^=0.54); atrial myocytes with higher levels of PAM staining generally had higher levels of proANP staining. In *Pam*^Myh6-cKO/cKO^ atrial myocytes, PAM staining was undetectable, but proANP levels again varied widely (**Fig. 4D**, gray boxes).

Atrial myocytes prepared from post-natal day 4 *Pam*^Myh6-cKO/cKO^ pups were transduced with lentiviruses encoding PAM1 or PAM1/Met314Ile at the time of plating and examined 7 days later (**Fig. 4C**). PAM and proANP staining intensities were quantified in *Pam*^Myh6-cKO/cKO^ cells expressing PAM1 or PAM1/Met314Ile (**Fig. 4E**). Strikingly, as levels of PAM1 expression rose, levels of proANP staining generally rose; the slope of this regression line was not as steep as the slope observed in control cells (solid black line from **Fig. 4D**), but levels of proANP and PAM were highly correlated (R^2^=0.51, blue squares). There was a wide range of expression of PAM from the Cre-activated lentivirus, as expected from similar studies with isolated β-cells and Cre-activated GFP expression (44). Analyses using a different pair of antibodies gave similar results (**Fig. S4**). Despite developing in the absence of PAM, the ability of atrial myocytes to store proANP was partially restored by expression of PAM.

Strikingly, in *Pam*^Myh6-cKO/cKO^ myocytes expressing PAM1/Met314Ile, a similar correlation of proANP and PAM expression was observed (**Fig. 4E**) (R^2^=0.60, orange diamonds). Based on this metric, the effects of PAM1 and PAM1/Met314Ile on the ability of atrial myocytes to store proANP were indistinguishable. Although the ability of PAM1 to increase proANP storage in atrial myocytes required high levels of PAM expression, it did not depend on its monooxygenase activity.

### Increased basal secretion and turnover of proANP by *Pam* ^Myh6-cKO/cKO^ atrial myocytes

To assess the possibility that the absence of PAM might trigger ER stress, we evaluated the levels of several stress responsive cardiomyocyte proteins and transcripts (**Fig. S5** and **Supplementary Table 2**) (45-47). The absence of PAM did not cause generalized ER stress or induce ER-phagy.

With a rise in *Nppa* transcript levels in *Pam*^Myh-cKO/cKO^ atria (3), a decrease in the synthesis of proANP or an increase in its degradation or secretion could contribute to the 3-fold decline in proANP levels in the adult atrium. Metabolic labeling was used to compare the ability of control and *Pam*^Myh-cKO/cKO^ atria to synthesize proANP (**Fig. 5A**). Total cell lysates and proANP immunoprecipitates were fractionated by SDS-PAGE and newly synthesized proANP was quantified by fluorography (**Fig. S6**). No significant change in proANP synthesis was observed in *Pam*^Myh-cKO/cKO^ atria when compared to control atria (**Fig. 5A**).

**Fig. 5.**
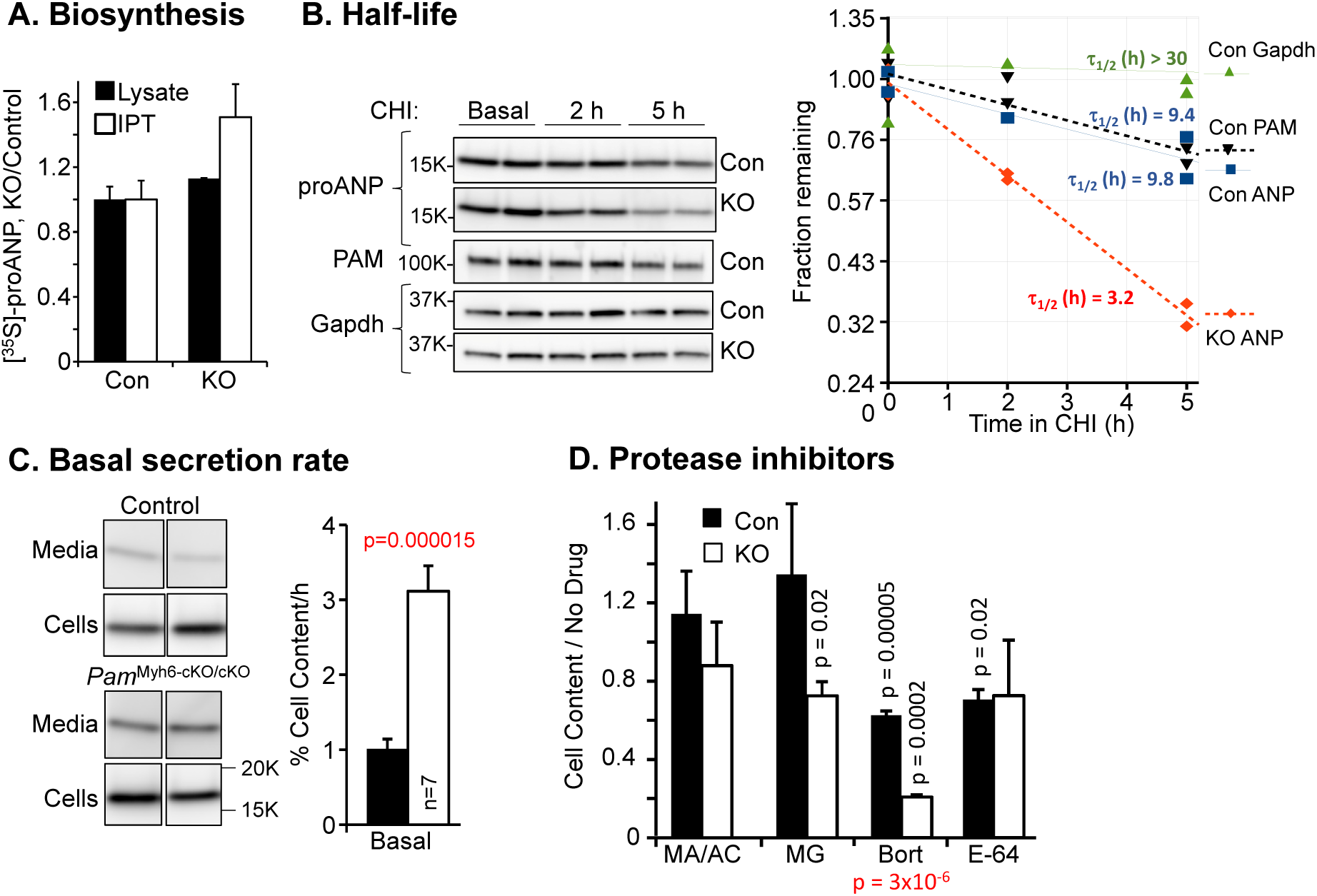
Evaluation of the role of PAM in proANP synthesis, half-life and secretion. **A. Biosynthesis**. Adult atrial tissue fragments (control and *Pam*^Myh6-cKO/cKO^) were incubated in medium containing [^35^S]Met/[^35^S]Cys for 15 min. RIPA soluble protein (Lysate) and proANP immunoprecipitates (IPT) were fractionated by SDS-PAGE and newly synthesized proANP was quantified by fluorography; proANP is the major 17 kDa protein in atrial extracts (15), allowing its direct quantification (**Fig. S6**). **B. Half-life.** The half-lives of proANP and PAM were determined using primary cultures of control and *Pam*^Myh6-cKO/cKO^ atrial myocytes. Duplicate cultures were incubated in medium containing 10 μM cycloheximide for the indicated periods of time and analyzed as described in **Methods**. Time course data for a single experiment are plotted (right side). **C. Basal secretion rate.** Myocytes prepared from control and *Pam*^Myh6-cKO/cKO^ pups were incubated in CSFM for 2 h; proANP levels in cell lysates and spent media were determined, with secretion rates expressed at % cell content secreted per h. **D. Protease inhibitors**. Cultures treated with methylamine/NH_4_Cl for 2-12 h or with MG-132 for 2-5 h, Bortezomib or E64 for 2 h were extracted for SDS-PAGE analysis of proANP levels; for each genotype, inhibitor treated lysates were compared to corresponding control lysates. For panels **C.** and **D.**, p values in red report a genotype-specific effect of a drug (control vs. KO) while p values in black report the effect of a specific drug on cultures of the indicated genotype.

We next used cycloheximide to inhibit protein synthesis and determine the effect of genotype on the half-life of proANP. A genotype-specific difference in the stability of proANP was apparent by 2 h (**Fig. 5B**). The half-life of proANP in *Pam*^Myh6-cKO/cKO^ myocytes (5.5 ± 0.9 h; n = 4) was approximately half that of proANP in control myocytes (11.7 ± 1.3 h; n = 5; p < 0.004). For comparison, the half-life of PAM in control myocytes was 7.8 ± 1.1 h. As expected, the half-life of Gapdh in both control and KO cultures was substantially greater than 30 h (48).

The decreased half-life of proANP in *Pam*^Myh6-cKO/cKO^ myocytes could reflect an increase in secretion and in degradation. To evaluate secretion rates, cultures were incubated in complete serum free medium for 2 h; proANP levels in cell lysates and media were evaluated by Western blot and secretion rates (relative to cell content) were determined (**Fig. 5C**). *In vivo*, proANP is cleaved at the moment of secretion by Corin, an endoprotease located on the sarcolemma (4). When maintained under the culture conditions used in this study, cardiomyocytes secrete primarily intact proANP (49, 50). The proANP band detected in spent media was indistinguishable from the proANP band in cell lysates, with little evidence of significant cleavage. While control myocytes released 1.0 ± 0.1% of their content of proANP per hour, *Pam*^Myh6-cKO/cKO^ myocytes released 3.1 ± 0.3% of their content of proANP per hour (**Fig. 5C**). This increase in proANP basal secretion would contribute to its decreased half-life in *Pam*^Myh6-cKO/cKO^ myocyte cell extracts. The decreased levels of mature ANP observed in the sera of *Pam*^Myh6-cKO/cKO^ mice may reflect a lack of proANP cleavage (3).

To determine whether increased degradation also contributed to the more rapid disappearance of proANP from *Pam*^Myh6-cKO/cKO^ myocytes, cultures of both genotypes were treated with methylamine/ammonium chloride to raise luminal pH and inhibit lysosomal degradation, with MG132 or Bortezomib to inhibit proteasomal degradation or with E64 to inhibit calpain cleavage (**Fig. 5D**) (51). None of these treatments increased the proANP content of control or *Pam*^Myh6-cKO/cKO^ atrial myocytes. Instead, MG132 reduced the proANP content of *Pam*^Myh6-cKO/cKO^ atrial myocytes and Bortezomib reduced the proANP content of both control and *Pam*^Myh6-cKO/cKO^ atrial myocytes. While our ultrastructural studies revealed a decrease in the diameter of Golgi-localized lysosomes in *Pam*^Myh6-cKO/cKO^ vs control atria (**Fig. 2K**) and the protease inhibitors tested failed to increase proANP levels in *Pam*^Myh6-cKO/cKO^ myocytes, increased proANP degradation may contribute to the decreased proANP content of *Pam*^Myh6-cKO/cKO^ myocytes.

### Basal secretion of proANP by control and *Pam*^Myh6-cKO/cKO^ myocytes differs in its sensitivity to BFA

We employed several well characterized pharmacological tools to explore the mechanism underlying the increased basal rate at which proANP is secreted by *Pam*^Myh6-cKO/cKO^ atrial myocytes. Based on the literature, most of the proANP released basally by control atrial myocytes is newly synthesized (13, 52, 53). Consistent with these earlier studies, cycloheximide treatment reduced basal proANP secretion by control atrial myocytes to 42 ± 13 % of the solvent control (**Fig. 6 A, B**). Cycloheximide had a smaller effect on the elevated basal secretion of proANP by *Pam*^Myh6-cKO/cKO^ myocytes, reducing it to 63 ± 9 % of the solvent control (**Fig. 6A, B**; n = 9, p = 0.0002) and suggesting a greater contribution of older proANP to basal secretion in atrial myocytes lacking PAM. The effects of cycloheximide on cell content of proANP over the course of this 2 h incubation were consistent with the half-lives reported in **Fig. 5B**, with a significant decline detectable in *Pam*^Myh6-cKO/cKO^ myocytes but not in control myocytes (**Fig. 6C**).

**Fig. 6.**
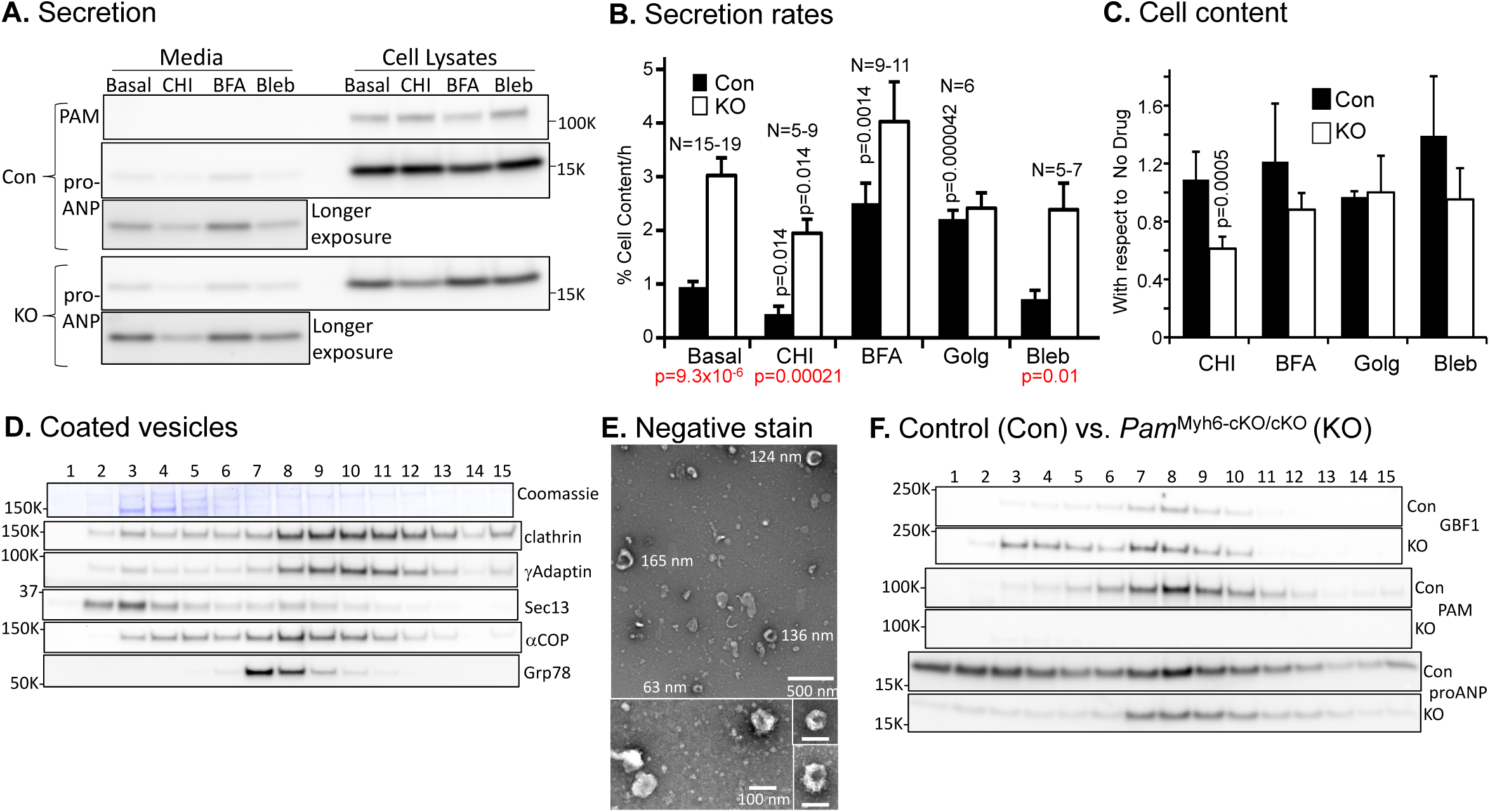
Pharmacological manipulation of the secretory pathway. **A. Secretion** and **B. Secretion Rate**. The effects of cycloheximide (10 μM, with a 1 h pre-treatment), Brefeldin A (BFA, 7.1 μM), and Golgicide A (10 μM) on the secretion of proANP by control (Con) and *Pam*^Myh6-cKO/cKO^ (KO) myocytes were determined. In the example shown, the litter used to prepare cultures contained 1 Con and 5 KO pups; media and cell extracts were analyzed simultaneously. Secretion rates are expressed as % proANP cell content released/h; n values for each comparison are shown. Statistics: p values for drug vs. control of the same genotype are shown in black and p values for genotype-specific drug effects are shown in red. **C. Cell content.** Drug effects on cell content of proANP (proANP/Gapdh) in control and KO cultures are shown. **D. Coated vesicles**. A coated vesicle enriched fraction prepared from adult control atria was subjected to sucrose density centrifugation. The Coomassie-stained membrane is shown above Western blot analyses for the indicated marker proteins; similar results were obtained in three additional experiments. **E. Negative stain.** Aliquots of Fraction 8 were spotted onto a grid, rinsed and negatively stained. Representative images are shown, with diameters (nm) of selected vesicles indicated. **F. Control and KO coated vesicles.** Coated vesicles prepared simultaneously from Con and KO atria were subjected to sucrose gradient fractionation. Expression of GBF1, PAM and proANP was evaluated in aliquots representing equal amounts of input protein.

The movement of cargo proteins from their site of synthesis in the ER through the Golgi complex and into secretory granules involves vesicular trafficking controlled by several Arf family small GTP binding proteins. BFA, an inhibitor of the Golgi-localized Arf GEFs (10-12), stimulates the basal secretion of proANP (13, 14). In contrast, BFA inhibits aspects of the basal and constitutive-like secretion of insulin, gastrin and ACTH (8, 9) (**Fig. S7**). As expected, BFA produced a 2.5 ± 0.4-fold increase in proANP secretion by control atrial myocytes (**Fig. 6A, B**; n = 11, p = 0.0014). Strikingly, BFA had no effect on the basal secretion of proANP by *Pam*^Myh6-cKO/cKO^ myocytes (**Fig. 6A, B**; n =9, p = 0.21). Cell content of proANP was not affected by this 2 h BFA treatment in cultures of either genotype (**Fig. 6C**).

Golgicide A inhibits the activity of Golgi BFA resistance factor 1 (GBF1), the Arf GEF localized to the *cis*-Golgi area, without affecting the activity of BFA-inhibited GEF1 (BIG1) or BFA-inhibited GEF2 (BIG2), the Arf GEFs localized to later parts of the secretory pathway (10, 11). The actions of Golgicide A on basal proANP secretion by control and *Pam*^Myh6-cko/cko^ myocytes mimicked those of BFA, with a stimulatory response by control myocytes and no effect on *Pam*^Myh6-cko/cko^ myocytes (**Fig. 6A, B**). Corticotrope tumor cells were used to verify the ability of BFA and Golgicide to inhibit specific aspects of the basal, constitutive-like and regulated secretion of prohormone convertase 1 and proopiomelanocortin products (**Fig. S7**). The fact that basal proANP secretion by atrial myocytes lacking PAM resembled basal proANP secretion by Golgicide A-treated control myocytes suggested that inhibition of GBF1 prevented PAM from carrying out its normal role in the secretion of proANP by atrial myocytes.

Blebbistatin, a myosin II-specific inhibitor (54) blocks tropomyosin 4.2 dependent ER to Golgi trafficking in mouse embryonic fibroblasts (55) and BFA-induced retrograde *cis*-Golgi to ER trafficking in pheochromocytoma cells (56). Blebbistatin had no effect on the rate at which proANP was secreted by either control or *Pam*^Myh6-cKO/cKO^ myocytes (**Fig. 6A, B**).

### PAM has a rate-limiting role in atrial granule biogenesis

The dramatic drop in granule number in *Pam*^Myh6-cKO/cKO^ atria suggested a role for PAM in granule biogenesis; proANP accounts for over 95% of the soluble protein in atrial granules (16, 52). Evidence for a direct interaction of PAM with proANP comes from the expression of fluorescently tagged PAM in cardiomyocytes (18), suggesting that when present at sufficiently high levels, PAM might interact with proANP, acting as a chaperone or transporter. Consistent with this idea, loss of a single *Pam* allele reduced proANP levels (**Fig.3A**) and rescue of proANP storage in atrial myocytes lacking PAM was dependent on the level of PAM expression (**Fig. 4E**). Taken together, the 1:30 molar ratio of PAM:proANP in rat atrial granule membranes (16) and the dose-dependent effects of PAM expression suggest that PAM is present in limiting amounts.

The ability of catalytically inactive PAM to restore proANP storage supports this hypothesis, suggesting that recycling of PAM is required for normal granule formation. The recruitment of CopI coats to membranes of the *cis*-Golgi is blocked by depletion or inhibition of GBF1 (10). If inhibiting GBF1 with Golgicide blocked the return of PAM from the *cis*-Golgi to the ER, proANP traversing the control cell secretory pathway might be susceptible to basal release, as in *Pam*^Myh6-cKO/cKO^ myocytes.

To assess this hypothesis, we prepared a coated vesicle enriched fraction from adult mouse atria (**Fig. 6D**). Based on the localization of clathrin, γ-Adaptin, Sec13 (which dissociates from CopII vesicles) and α-Cop (a marker for CopI vesicles), fractions 8-12 were enriched in coated vesicles. Grp78, a luminal ER chaperone that is retrieved via the KDEL receptor and CopI vesicles, was more enriched in lighter fractions. Negative staining of sucrose density gradient Fraction 8 confirmed the presence of coated vesicles, many of which resembled CopI coated vesicles (**Fig. 6E**). With the small amounts of material available, we did not attempt to separate CopI and CopII coated vesicles.

Based on our hypothesis, CopII coated vesicles should contain newly synthesized PAM and newly synthesized proANP while CopI coated vesicles should contain PAM being recycled to the ER, but not proANP. PAM was recovered in coated vesicle enriched fractions prepared from control atria (**Fig. 6F**). The sonication step used to prepare coated vesicles would be expected to completely disrupt atrial granules, releasing soluble content proteins like proANP, and only a small fraction of the proANP present in each lysate was recovered in the high-speed pellet. Consistent with the greatly reduced number of granules in *Pam*^Myh6-cKO/cKO^ atria, very little proANP was recovered from the top of this gradient; proANP was recovered from the coated vesicle enriched fractions of both control and KO atria (**Fig. 6F**).

Levels of the mRNAs encoding GBF1, the Arf proteins and the coatomer subunits (**Table S2**) and levels of GBF1, Sec13 and Sar1B protein (**Fig. S8A** and **B**) did not differ in wildtype and *Pam*^Myh6-cKO/cKO^ atria. Based on subcellular fractionation, GBF1 was enriched in the P3 and P4 fractions of control and *Pam*^Myh6-cKO/cKO^ atria (**Fig. S8C**), co-localizing with PAM during sucrose gradient fractionation of control P3 and P4 fractions (**Fig. S8D**). GBF1 was recovered from coated vesicle enriched fractions of control and *Pam*^Myh6-cKO/cKO^ atria. The fact that its distribution differed, with more GBF1 in the lighter fractions of *Pam*^Myh6-cKO/cKO^ atria, supports the hypothesis that PAM affects ER/Golgi vesicular trafficking.

### PAM facilitates proANP storage in atrial granules by altering its pre-Golgi trafficking

We used HEK293 cells, which lack secretory granules and any of the features unique to cardiomyocytes, to test the hypothesis that PAM alters the trafficking of proANP at an early stage in the secretory pathway. ProANP synthesis requires formation of an intrachain disulfide bond, but does not require N-glycosylation, endoproteolytic cleavage or extensive O-glycosylation, making it difficult to track its movement through the secretory pathway (57). proANP-Emerald (58, 59) was transiently expressed in HEK293 cells and in HEK293 cells stably expressing PAM1 (PAM/HEK) at levels similar to those observed in adult mouse atrium (**Fig. S9A**). proNPY-GFP (60), a control for cargo efficiently targeted to secretory granules in neuroendocrine cells, was also transiently expressed in HEK293 and PAM/HEK cells (**Fig. S9B, S9C**).

When expressed in HEK293 cells, proANP-Emerald and proNPY-GFP were both diffusely distributed (**Fig. 7A**). In contrast, when expressed in PAM/HEK cells, proANP-Emerald was often condensed in what appeared to be tubulovesicular clusters located in the perinuclear region (**Fig. 7A**). In cells expressing lower levels of proANP-Emerald, a more diffuse accumulation of proANP-Emerald was observed in the perinuclear region (**Fig. 7A**). The localization of transiently expressed proNPY-GFP was less affected by the presence of PAM, but its accumulation in the perinuclear region of PAM/HEK cells was observed (**Fig. 7A**). Epifluorescence images of fixed HEK293 and PAM/HEK cells expressing proANP-Emerald or proNPY-GFP were categorized as exhibiting a diffuse, peri-nuclear or condensed pattern of immunofluorescence (**Fig. 7B**). The presence of PAM had different effects on the localization of proANP-Emerald and proNPY-GFP; while condensed peri-nuclear staining was observed with proANP-Emerald, diffuse peri-nuclear staining was observed with proNPY-GFP (**Fig. 7B**).

**Fig. 7.**
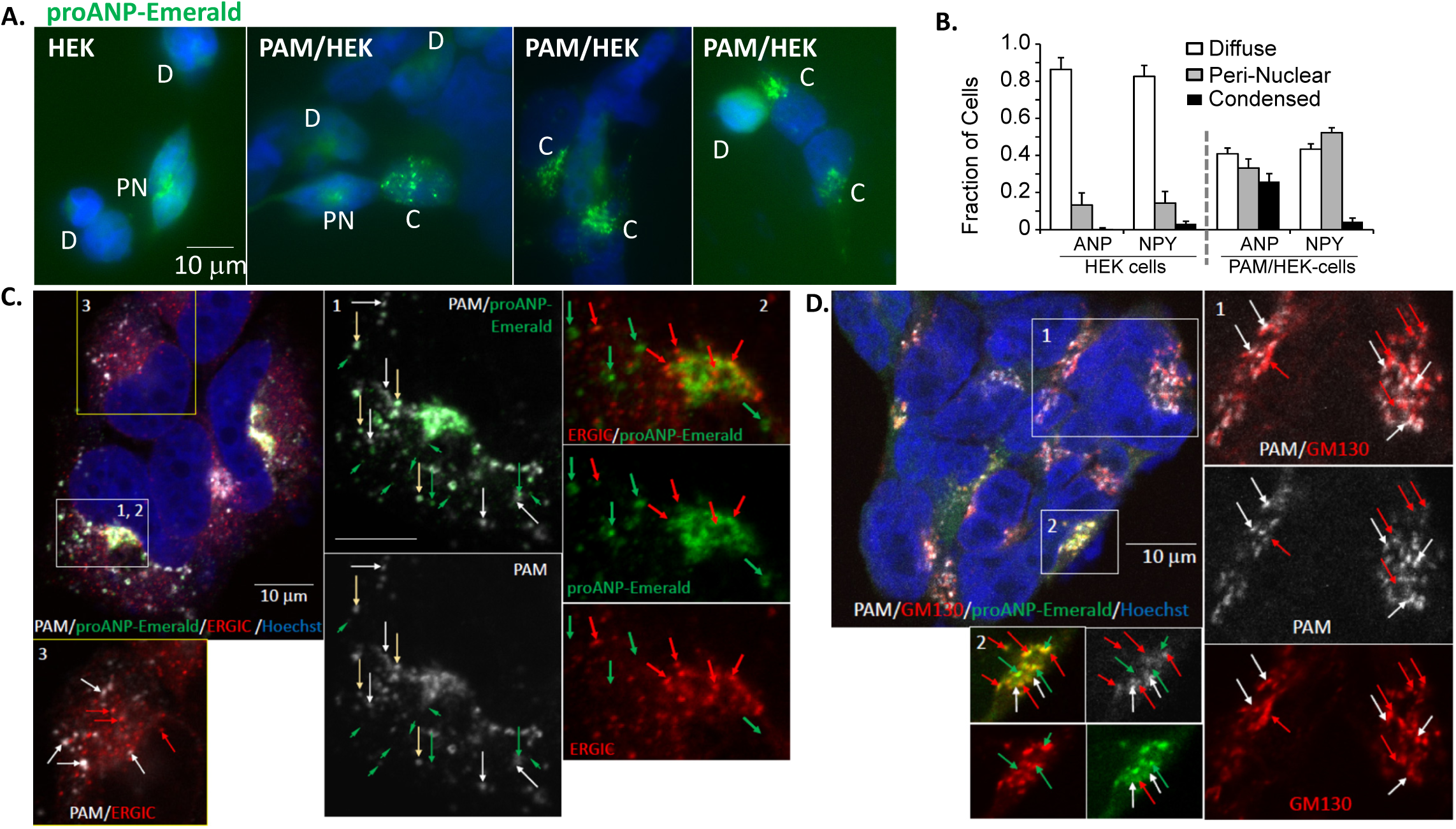
Expression of proANP-Emerald and proNPY-GFP in HEK cells and PAM/HEK cells. Sparsely plated HEK cells and PAM/HEK cells transiently transfected the previous day with vectors encoding proANP-Emerald or proNPY-GFP (not shown) were fixed; nuclei were visualized using Hoechst stain and epifluorescence images of the transiently expressed proteins were collected using a 40X objective. Coverslips were scanned systematically and cells expressing a fluorescently-tagged protein were imaged under standard conditions. Fluorescent protein localization was categorized as diffuse (D), peri-nuclear (PN) or condensed (C) by a blinded observer. **A.** Examples illustrating the three staining categories are shown for proANP-Emerald expressed in HEK or PAM/HEK cells; scale bar, 10 μm for all images. **B.** Quantitative data for each cell type and vector are shown; the number of cells and number of groups averaged were HEK-ANP (236; 4), HEK-NPY (932; 7), PAM/HEK-ANP (514; 5), PAM/HEK-NPY (222; 5). Based on a two-way ANOVA, the behavior of the two cell lines and the two fluorescently tagged proteins differed (p < 0.0001 for all comparisons). **C.** and **D.** Confocal images were obtained for PAM/HEK cells expressing proANP-Emerald (green) and stained for PAM (gray) and ERGIC-53 (red) (**C**) or for PAM (gray) and GM130 (red) (**D**). Numbered insets are shown at higher magnification with single colors or two colors, as indicated.

Confocal images of PAM/HEK cells expressing proANP-Emerald revealed extensive overlap of PAM and proANP-Emerald in reticular structures and in puncta (yellow arrows) (**Fig. 7C, Inset 1**). Small puncta containing proANP (short green arrows) or PAM (white arrows) were also seen (**Fig. 7C, Inset 1**). The localization of proANP and PAM was compared to that of ERGIC-53, a marker for ER exit sites (ERESs) and the tubulovesicular clusters responsible for the bidirectional traffic that connects the ER to the *cis*-Golgi (ER/Golgi intermediate compartment) (**Fig. 7C, Insets 2** and **3**) (61, 62). ERGIC-positive puncta (red arrows) were often localized adjacent to proANP-Emerald- and PAM-positive tubulovesicular structures (**Insets 2** and **3**, respectively), but neither proANP-Emerald nor PAM accumulated in the ERGIC-53 positive puncta. Extensive overlap was observed when PAM/HEK cells expressing proANP-Emerald were stained for PAM and GM130 (**Fig. 7D**). Staining for PAM (**Inset 1**, white arrows) and for GM130 (**Inset 1**, red arrows) was frequently overlapping or immediately adjacent. Staining for proANP and GM130 (**Fig. 7D, Inset 2**) was often coincident. Taken together, our data indicate that expression of PAM facilitates the accumulation of proANP in the *cis*-Golgi region and that proANP does not accumulate at ERESs.

## DISCUSSION

### PAM plays a role in basal proANP secretion and granule formation

Eliminating *Pam* expression in atrial myocytes resulted in a 3-fold drop in proANP levels in the atrium, a 20-fold decline in granule cytoplasmic volume fraction and a 3-fold increase in proANP basal secretion. While the decrease in proANP levels can be accounted for in part by the increase in basal secretion, the almost complete absence of atrial granules suggested a specific role for PAM in granule formation or stability. PAM expression in the atrium exceeds levels in all other tissues; since the natriuretic peptides are not amidated, it was proposed that PAM facilitates their condensation and storage, as observed for secretogranins (15, 16). Our data establish a non-catalytic role for PAM in the atrium and localize its actions to early stages of the secretory pathway (**Fig. 8**).

**Fig. 8.**
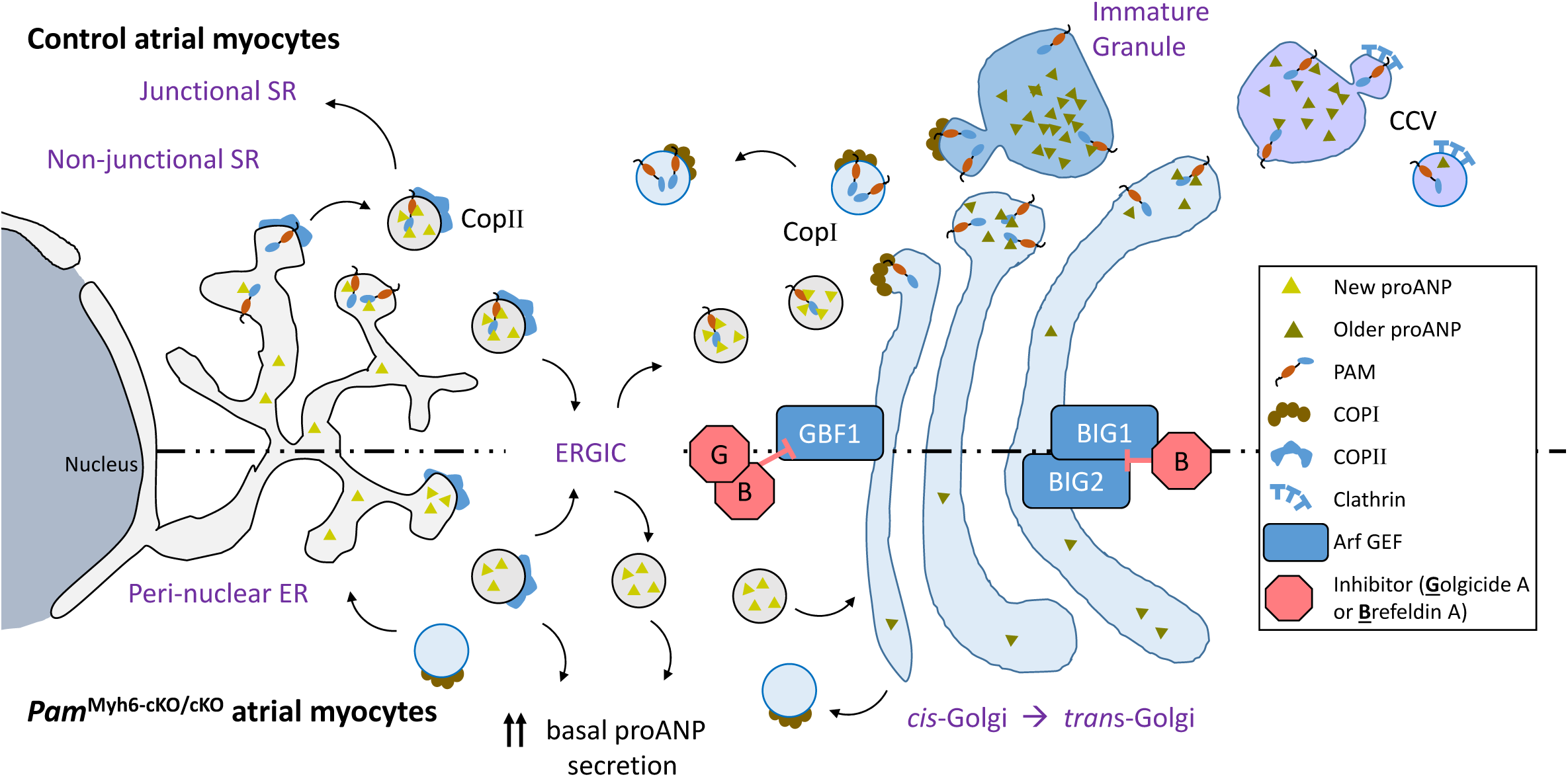
Working Model. Key differences in the trafficking of newly synthesized proANP from its site of synthesis in the peri-nuclear ER through the ER/Golgi Intermediate Compartment (ERGIC) and into the *cis*-Golgi and immature secretory granules in Control (above the dashed line) and *Pam*^Myh6-cKO/cKO^ atrial myocytes are highlighted.

Demonstration of a direct interaction between PAM and proANP will require additional experimentation, but multiple observations support the proposal. A conserved pair of adjacent acidic residues near the N-terminus of proANP plays an essential role in its ability to bind Ca^2+^, aggregate and enter the regulated secretory pathway (63). These same residues play an essential role in the ability of proANP to interact with PAM in transfected atrial myocytes (18, 58). PAM and proANP together account for 95% of the protein in atrial granule membranes (16) and the effects of PAM on proANP storage were dose dependent and selective. Natriuretic peptide levels in the atria of mice with a single *Pam* allele were reduced, but not to the levels observed in *Pam*^Myh6-cKO/cKO^ mice; CPE levels were unaltered. Expression of exogenous PAM increased proANP storage in *Pam*^Myh6-cKO/cKO^ atrial myocytes in a dose dependent manner that did not require its monooxygenase activity. When transiently expressed in cells which lack secretory granules and express PAM at low levels, proANP-Emerald was diffusely distributed; in lines stably expressing PAM at levels observed in the atrium, proANP-Emerald and PAM were largely localized near GM130-positive, perinuclear structures.

Taken together, these diverse observations support a working model built around a low affinity, direct interaction between the luminal domains of PAM and proANP (**Fig. 8**). The model incorporates a direct interaction of PAM with newly synthesized proANP, facilitating proANP delivery to the *cis*-Golgi and to immature granules and preventing its basal secretion (**Fig. 8**). The ability of Golgicide A, a specific inhibitor of GBF1, the Arf-GEF localized to the *cis*-Golgi, to stimulate proANP secretion by control atrial myocytes without affecting proANP secretion by *Pam*^Myh6-cKO/cKO^ atrial myocytes focused our attention on the mechanism through which PAM limited basal proANP secretion. Studies on the roles of the luminal and cytosolic domains of PAM in its trafficking through the biosynthetic and endocytic compartments of corticotrope tumor cells guided our interpretation of its role in myocytes.

### Basal secretion of proANP

Studies using intact atria and primary rodent atrial myocyte cultures established the key features of basal proANP release. Unlike most peptide hormones and neuropeptides, much of the proANP secreted basally is recently synthesized (13, 53), suggesting that it might not have traversed the Golgi complex. While increased intracellular calcium stimulates the secretion of most peptide hormones and neuropeptides (8, 9) (**Fig. S7**), it inhibits the basal secretion of natriuretic peptides (5, 6). Unlike the stimulated secretion of proANP, its basal secretion is not affected by pertussis toxin or by monensin. BFA treatment of cannulated atria or cultured atrial myocytes increases basal proANP release (13, 14). Using primary cardiomyocyte cultures, we confirmed the ability of BFA to increase basal proANP release and demonstrated the ability of Golgicide A to do the same. Based on these observations, basal proANP secretion does not require Arf-activated vesicular trafficking, suggesting that it comes largely from a source that precedes the *cis*-Golgi.

Unique features of the ER/SR in atrial myocytes may contribute to the unusual features of proANP secretion. Transverse tubules, which are prevalent in ventricular myocytes, are rare in atrial myocytes (64). Instead, junctional SR underlies the sarcolemma, connected to the perinuclear ER by non-junctional SR. Calsequestrin, a soluble calcium binding protein, moves from its site of synthesis in the perinuclear ER to the lumen of the junctional SR, where it plays a role in calcium release; this process is inhibited by disruption of COPII-mediated vesicular budding, but not by disruption of intra-Golgi trafficking (65). The machinery that allows calsequestrin to travel from its site of synthesis to the junctional SR could afford a pathway through which newly synthesized proANP is basally secreted without traversing the Golgi complex.

Some membrane proteins exit the ER in non-COPII vesicles or by-pass the Golgi complex, yet still reach the plasma membrane (66). The cystic fibrosis transmembrane conductance regulator (CFTR) exits the ER in a CopII-dependent manner, but by-passes the Golgi complex, accumulating in a peri-centriolar intermediate compartment (66). In neurons, a significant fraction of the forward trafficking from ER to dendritic recycling endosomes and the plasma membrane occurs even when the Golgi apparatus has been disrupted by BFA and many surface receptors have immature N-linked glycans, consistent with Golgi bypass (61). A Golgi by-pass pathway would allow production of proANP, which does not require N-glycosylation, is not extensively O-glycosylated and is cleaved only at the time of secretion (57).

### PAM protects newly synthesized proANP from basal release and delivers proANP to nascent granules

The exit of newly synthesized proteins from the ER involves bulk flow and active sorting. Soluble cargo proteins like proANP can enter CopII vesicles by diffusing into a budding zone, or can be concentrated in CopII vesicles through interactions with p24 family proteins or general cargo receptors like ERGIC-53, a lectin with CopII sorting signals (67). The cytosolic domain of PAM has highly conserved diacidic and dihydrophobic motifs resembling those that allow the vesicular stomatitis virus glycoprotein and ERGIC-53, respectively, to bind to CopII coat proteins (67). While the ER is a single continuous structure, different functions are carried out in distinct zones. Cargo often exits the ER at ribosome-free subdomains [ER exit sites (ERESs) or transitional ER], where COPII coated vesicles are concentrated (67). In vertebrate cells, ERESs are distributed across the ER, with many far removed from the Golgi complex. The vesicular tubular clusters that comprise the ER-Golgi intermediate compartment (ERGIC) receive cargo from ER exit sites, and link it to the *cis*-Golgi (68) (**Fig.8**).

The rate at which proANP was basally secreted by atrial myocytes lacking PAM equalled the rate at which proANP was secreted by BFA-treated atrial myocytes expressing PAM, suggesting that PAM protected newly synthesized proANP from basal secretion. Although the pathway through which BFA stimulates the basal secretion of proANP has not been elucidated, the fact that it does so only in the presence of PAM suggested the involvement of altered PAM trafficking. The ability of Golgicide A to mimic this effect of BFA identified the *cis-*Golgi as the key site for this effect. With proANP levels exceeding PAM levels, PAM must be recycled to the ER after accompanying proANP to the *cis*-Golgi and perhaps to nascent secretory granules (**Fig. 8**). Blocking the return of PAM to the ER would leave newly synthesized proANP susceptible to secretion via an early BFA-insensitive pathway. A study of procathepsin secretion after its removal from immature secretory granules in β-cells revealed a stimulatory effect of BFA on its release from recycling endosomes (7).

There is a precedent for CopI-dependent return of cargo receptors to the ER. Yeast Erv29p and its *C. elegans* homolog, SFT-4/Surf4 (Surfeit locus protein 4 homolog), are ER-localized membrane proteins required for the efficient CopII-mediated export of specific soluble cargo proteins; these cargo receptors are returned to the ER in CopI coated vesicles (69). TANGO1 (Transport and Golgi organization protein 1), a Type 1 ER-localized membrane protein, delivers procollagen to departing COPII vesicles, but remains in the ER (67). Our data suggest that the actions of PAM resemble those of Erv29p and SFT-4.

The cytosolic domain of PAM interacts with the μ1A subunit of the AP-1 complex (70), with Rho GEFs that activate Rac1, RhoG and RhoA (71) and with actin (72). In corticotrope tumor cells, PAM enters immature secretory granules, produces amidated products and is removed in an AP-1 dependent process. The ability of the cytosolic domain of PAM to interact with μ1A plays an essential role in granule maturation, and the endocytic trafficking of PAM is guided by the reversible phosphorylation of multiple Ser/Thr residues in its cytosolic domain. The μ1A subunit most closely resembles the δCOP subunit of CopI (73). PAM was identified in atrial coated vesicle enriched fractions. Subcellular fractionation revealed enrichment of GBF1 in the subcellular fractions that contain PAM; αCop, GBF1 and PAM were similarly localized following sucrose density gradient fractionation of atrial coated vesicles. The fact that PAM expressed in cells lacking secretory granules localized to the *cis*-Golgi region and altered the localization of transiently expressed proANP-Emerald is consistent with a role for PAM in the early secretory pathway, preceding the Golgi complex. The effects of PAM on soluble cargo trafficking are not limited to proANP. High levels of PAM expression in HEK cells led to proNPY accumulation in the perinuclear region. In AtT-20 corticotrope tumor cells, PAM expression leads to POMC accumulation in the *trans*-most cisternae of the Golgi complex, limiting its cleavage and storage of its product peptides in secretory granules (74).

Left atrial function is an independent predictor of adverse cardiac events in the general population and in hypertension and heart failure patients (75) and atrial failure, in the absence of ventricular or valvular abnormalities, can cause heart failure (76). The current understanding of protein trafficking, signaling and cytoskeletal control in atrial myocytes is limited. In *Pam*^Myh6-cKO/cKO^ atrial myocytes, fewer Golgi complexes were found near the sarcolemma. Left atrial myocytes in tissue from atrial fibrillation patients exhibited Golgi fragmentation, with an increase in smaller fragments localized lateral to the nucleus (25). Understanding how rhythmic contraction, a known regulator of the atrial myocyte cytoskeleton (77), affects the protein trafficking network in which PAM and proANP function may reveal new ways in which to manipulate the system.

## METHODS

PAM^cKO^ mice are available from The Jackson Laboratory as JAX#034076. All work with mice was approved by the University of Connecticut Health Center Institutional Animal Care and Use Committee. Detailed Methods appear in Supplementary Material.

## ACKNOWLEDGEMENTS

This work was supported by a grant from the National Institutes of Health (DK032948 to REM) and support from the Rodney and Janice Reynolds Endowment (to BAE), the Daniel Schwartzberg Fund (to REM and BAE) and the Finska Läkaresällskapet and Perklén Foundation (to NB). Maya Yankova’s expertise and the UConn Health Central EM Facility made our initial analysis of knockout atria and our negative stain analysis of atrial coated vesicles possible. We thank the Electron Microscopy Unit of the Institute of Biotechnology, University of Helsinki for providing laboratory facilities. We thank Dr. Chris Glembotski (San Diego State University) for his generous supply of rabbit antiserum to the N-terminal region of proANP. We thank Drs. Stephen King (UCHC) and Yi-Chun Chen (University of British Columbia) for their comments.

## METHODS

### Mouse husbandry

*Pam*^cKO/+^ mice are available from The Jackson Laboratory as JAX#034076 (1). JAX011038-B6.FVB-Tg(Myh6-cre)2182Mds/J mice, which are also on a C57BL/6J background (The Jackson Laboratory), were maintained as hemizygotes. Genotyping was performed from ear or tail clips as described (1). On average, breeding a homozygous *Pam*^cKO/cKO^ mouse with a hemizygous for Myh6-Cre recombinase mouse yielded an equal number of *Pam*^Myh6-cKO/cKO^ (cardiomyocyte specific *Pam* knockout) and *Pam*^cKO/cKO^ (control) mice; PAM^cKO/+^ and pure C57BL/6J mice also served as controls.

### SDS-PAGE analysis of tissue lysates

Atria (left and right) from adult (41-128 days old) male and female C57Bl6/J mice were rinsed in PBS to remove most of the blood and were then sonicated in 100 μl ice cold RIPA buffer (Cell Signaling Technologies, #9806) containing 0.3 mg/ml phenylmethylsulfonyl fluoride. After incubation on ice for 10 min, insoluble material was removed by centrifugation at 17,200 xg for 10 min and the concentration of protein in the soluble fraction was measured using the bicinchoninic acid assay with bovine serum albumin as the standard (Thermo). Samples from control mice (wild type, *Pam*^cKO/cKO^, *Pam*^cKO/+^ and Myh6/+; n = 5), heterozygous mice (*Pam*^Myh6-cKO/+^; n= 5) and knockout mice (*Pam*^Myh6-cKO/cKO^; n = 8) were analyzed. Since no differences were observed between lysates from males and females, data were grouped by genotype. Depending on the proteins to be analyzed, samples (20 μg protein) were analyzed on 18-lane or 12+2 lane 12% polyacrylamide or 4-15% polyacrylamide Bio-Rad Criterion TGX gels (BioRad). Following transfer to a PVDF membrane, levels of PAM, subcellular marker proteins, cargo proteins and PAM interactors were quantified using HRP-tagged secondary antibodies (Jackson ImmunoResearch) and SuperSignal West Pico chemiluminescent substrate (Pierce). Images were captured using GeneTools (SynGene, Frederick MD) and signal intensities were quantified from non-saturated images. Additional lysates were prepared by sonication of atrial tissue or cell pellets into SDS-P lysis buffer (50 mM TrisHCl, 2% SDS, 130 mM NaCl, 5 mM EDTA, 50 mM NaF, 50 mM Na pyrophosphate, pH 7.6) containing 0.3 mg/ml freshly added phenylmethylsulfonyl fluoride and a protease inhibitor cocktail (P8340, Sigma); following sonication, samples were heated at 50°C or 95°C for 5 min and clarified by centrifugation at 16,000 xg for 20 min. Atrial lysates prepared for evaluation with the Cell Signaling Technologies ER Stress Antibody Sampler Kit (#9956, Cell Signaling Technologies) were prepared by sonication in 200 μl SDS-P buffer/(pair of atria); in addition to protease inhibitors, phosphatase inhibitors (vanadate and PHOSTOP) were added to the lysis buffer. Antibodies used for Westerns and immunocytochemistry are in **Supplemental Table 1**.

### Differential centrifugation and sucrose density gradient fractionation

Secretory granules account for less than 1% of the cytosolic volume of atrial myocytes (**Fig. 1E**), while myofilaments plus mitochondria account for approximately 75% (2). Published fractionation protocols (3-5) had to be modified to deal with the small amounts of atrial tissue available and the scarcity of granules. Atria (left and right) from adult mice (typically 4 mice; mixed sexes) were collected into ice cold 10 mM HEPES, 0.25 M sucrose, pH 7.4. Two scalpels were used to mince the chilled atria; tissue fragments were transferred into a Potter-Elvejelm homogenizer containing 0.5 ml homogenization buffer (10 mM HEPES, 50 mM EDTA, 0.25 M sucrose, pH 7.4 with a protease inhibitor cocktail, Sigma P8340). Following 12 up and down strokes with a motor driven Teflon pestle, samples were transferred into microfuge tubes for differential centrifugation. A crude nuclear pellet (P1) was obtained by centrifugation for 5 min at 1000 xg. A crude mitochondrial pellet (P2) was obtained by centrifugation of the supernatant for 10 min at 2000 xg. This supernatant was then centrifuged for 15 min at 32,000 xg in a TL100 rotor, yielding a crude granule fraction (P3). The final supernatant was centrifuged for 15 min at 435,000 xg, yielding a crude microsomal fraction (P4) and cytosol.

The P3 and P4 pellets were gently resuspended in 0.4 ml Homogenization Buffer using a hand held motor drive pestle; an aliquot of each was saved for analysis of the differential centrifugation paradigm and 350 μl was layered onto the top of a step gradient prepared from 10 mM HEPES, 25 mM EDTA, pH 7.4 containing varying amounts of sucrose: 0.25 ml of 2.0 M sucrose; 0.25 ml of 1.6 M sucrose; 0.50 ml of 1.4 M sucrose, 0.375 ml of 1.17 M sucrose, 0.375 ml of 0.93 M sucrose, 0.25 ml of 0.44 M sucrose. Gradients were centrifuged in a TLS55 rotor for 2 h at 120,000 xg_avg_. Samples were collected by sequentially removing fifteen 150 μl samples from the top of the tube; the final fraction was used to suspend any material that had pelleted. Aliquots of each fraction were subjected to Western blot analysis.

### Transmission electron microscopy and image analysis

Adult (2-2.5 month old) *Pam*^cKO/cKO^ (n=3) and *Pam*^Myh6-cKO/cKO^ (n=3) mice were perfused with saline (1) followed by 2.5% glutaraldehyde/2% formaldehyde (Electron Microscopy Sciences) in 0.1 M cacodylate buffer and post-fixed with the same fixative for 2 h. Cubes of left atrial tissue (∼1 mm^3^)were osmicated, dehydrated and embedded with random orientation in Epon. Sections were post-stained with uranyl acetate and lead citrate. Atrial tissue was systematically photographed at fixed intervals with a random start using a Jeol JEM-1400 electron microscope equipped with a Gatan Orius SC 1000B bottom mounted CCD-camera; a magnification of 600X was used for nuclear volume fractions and cell transection width and a magnification of 4000X was used for morphometric analysis of cell organelles. Two blocks from each animal were analyzed, with 20 photographs at each magnification for each block. Morphometric analysis was performed with the Stereology function in Microscopy Image Browser (6). Organelle diameters were analyzed with ImageJ. For analysis of Golgi complex transections, left atrial tissue was systematically scanned and Golgi complexes were photographed at 6000X to give 20 photographs/specimen.Irregular secretory granules were defined as granules with a clearly irregular outline or granules whose longest diameter exceeded its shortest diameter by ≥30%.

### Metabolic labeling and immunoprecipitation of proANP. Metabolic labeling experiments

Atria from two control and 2 *Pam*^Myh6-cKO/cKO^ post-natal day 30 male mice were diced and each atrium was incubated in 0.6 ml medium containing 2 mCi/ml [^35^S]-Met/[^35^S]-Cys (PerkinElmer EasyTagExpre35S35S, Waltham MA) in Met^-^ DMEM-F12 (Sigma P8340) for 15 min at 37C with rocking, rinsed for 30 sec in 2 ml ice-cold serum-free DMEM-F12, and homogenized with a glass-on-glass homogenizer into 0.1 ml RIPA buffer containing protease inhibitor cocktail P8340 (Sigma). Samples were centrifuged at 16,000 xg for 10 min, diluted into 0.5M KCl, 50 mM HEPES pH 7.3 (7), pre-cleared with Protein A resin for 1 h at 4C with rocking, incubated for 2 h with proANP antibody (8) at 4C, centrifuged at 16,000 xg for 10 min, and then incubated with Protein A resin for 1 h at 4C with rocking, rinsed twice in 20 mM Na TES, pH 7.4 containing 100 mM mannitol and 1.0% TX-100 (TMT) and fractionated by SDS gel electrophoresis. After impregnation with En3hance (Perkin Elmer), gels were exposed to film for up to 336h (9). The experiment was repeated with tissue from 1 control and 1 *Pam*^Myh6-cKO/cKO^ mouse with similar results.

### Preparation of primary mouse atrial cultures

Cultures were prepared from the atria of post-natal day 4 to 7 pups. Post-natal 3 to 4 pups were genotyped from tail clips and identified with paw tattoos, to allow pooling of tissue for preparation of control and *Pam*^Myh6-cKO/cKO^ cultures. In addition to control pups, cultures were prepared from the progeny of *Pam*^cKO/cKO^ x *Pam*^Myh6-cKO/cKO^ mice. Experiments were adjusted to accommodate the variable number of *Pam*^cKO/cKO^ (control) and *Pam*^Myh6-cKO/cKO^ (KO) pups available. The protocol used was modified from (10). Atria from P4-P7 pups were collected into serum-free medium and dissociated with a series of 5-10 min incubations at 37C with rocking in 1.5 ml trypsin (2.5 mg/ml Trypsin I-300 [ICN Biochemical 103140, Cleveland OH] in 5.4 mM KCl, 137 mM NaCl, 5.6 mM glucose, 20 mM Hepes, pH 7.35), with repeated pipetting with a P-1000 tip. Liberated cells were collected in 35 ml of growth medium (DMEM-F-12, 100 μg/ml kanamycin, 25 mM HEPES, 5% heat-inactivated Hyclone horse serum [GE Healthcare SH30074.03HI], 5% NuSerum [Corning 355500, Bedford MA], 0.5 mM ascorbate). Cells were filtered through a 70 μm filter, collected by centrifugation, incubated in 1 ml growth medium containing 1% Benzonase (Novagen, San Diego CA) for 10 min at room temperature, then another 10 min with 2 ml isotonic NH_4_Cl added [to lyse red blood cells]. Dissociated cells were collected by centrifugation, washed once in growth medium, preplated for 3 h into a 0.1% gelatin coated T25 flask to remove most fibroblasts, and plated in 100 μl on UV-treated, gelatin coated tissue culture plates, typically 1 atrium per well in a 96 well dish. Cells were fed daily for the first several days, then maintained in 200 μl medium until used for experiments (typically 3, 6 or 9 days).

To determine whether the differences observed between control and *Pam*^Myh6-cKO/cKO^ adult atria were mimicked in primary cultures, we normalized proANP levels in atrial cultures to Gapdh and to cardiac TnT (TnTc), to account for any differences in myocyte prevalence. As observed in adult tissue, proANP levels in *Pam*^Myh6-cKO/cKO^ atrial myocytes were reduced compared to control: the proANP/Gapdh ratio for KO vs. control cultures was 0.39 ± 0.04 (n = 8; p < 0.001) and the proANP/TnTc ratio was 0.35 ± 0.02 (n = 4; p < .001).

### Half-life experiments

Primary atrial cultures (wildtype, *Pam*^cKO/cKO^ [‘0-Cre’] and *Pam*^Myh6-cKO/cKO^ (KO)) were incubated without or with 10 μM cycloheximide (Calbiochem #239763) in growth medium for the indicated times (11), rinsed for 1 min in 37°C serum-free medium, extracted into SDS-P buffer (100 μl/well of a 96-well dish), processed as described above and then subjected to Western blot analysis for proANP, PAM and Gapdh. Images were quantified in the linear range using GeneTools (Syngene, Frederick MD). Data for cycloheximide-treated cultures were normalized to control cultures analyzed at the same time.

### Lentiviral manipulation of PAM expression in culture

cDNAs encoding active PAM1 and monooxygenase-inactive PAM1/Met314Ile (12, 13) were cloned into the G0655pFIV3.2CMVmcs vector (https://medicine.uiowa.edu/vectorcore/) and made into a high-titer feline immunodeficiency lentivirus at the University of Iowa Viral Vector Core (University of Iowa Health Care, Iowa City, IA). Crystallographic studies of PHM/Met314Ile, the soluble catalytic domain bearing the same mutation, revealed no overall change in the structure of this domain, with alterations localized to a short loop near the active site (13). Freshly dissociated and preplated atrial cells were spinoculated with 10^7^ TU virus in 100 μl cell suspension by centrifugation at 1200 xg for 3h, followed by a brief rinse in 2 ml growth medium (14, 15) and plated as above. Green fluorescent protein (GFP) and Cre-recombinase-GFP lentiviruses were also purchased from the University of Iowa Viral Vector Core. The integrity of each PAM lentivirus was determined after spinoculating pEAK Rapid cells (hEK derivative) (16) and assaying lysates for PHM and PAL activity (1) and protein integrity by Western blot analysis.

Elimination of PAM expression in *Pam*^cKO/cKO^ cultures was performed by spinoculation with the GFP (control) or Cre-GFP lentivirus. The time course over which protein expression changed was determined by Western blot analysis of SDS-lysates and by visualization of PAM and proANP in fixed cells by immunofluorescence at 3, 6 and 9 days after spinoculation. The PAM and proANP content of individual spinoculated *Pam*^Myh6-cKO/cKO^ cells expressing active PAM1 or enzymatically inactive PAM1/M314I was quantified by fixing and staining cultures 6 and 9 days after plating. Images of WT, KO and spinoculated cells were taken under identical conditions and levels of proANP and PAM staining were quantified, using Hoechst stain to distinguish single cells. PAM levels were quantified using a rabbit polyclonal antibody to the Exon A region of PAM-1 (JH629) and Alexa Fluor 488 donkey anti-rabbit IgG (A11029, Invitrogen, Thermo Fisher Scientific); ANP levels were quantified using a goat polyclonal antibody (Abcam ab190001) and Cy3-tagged donkey anti-goat IgG (705-166-147, Jackson ImmunoResearch Laboratories, Inc.). Quantification was also carried out using a mouse monoclonal antibody to PAM (6E6) with Alexa Fluor 488 goat anti-mouse IgG (A11029) and a rabbit antiserum to proANP (8) with Alexa Fluor 633 goat anti-rabbit IgG (A21052). Coverslips were systematically scanned for ANP-expressing cells; non-saturated images of ANP and PAM were acquired under fixed conditions. Quantification with FiJi (17) was accomplished by outlining ANP-positive cells and determining total integrated density for each fluorophore; background was assessed locally, adjusted for area and subtracted to obtain data for each fluorophore.

### Analysis of basal secretion by primary atrial myocytes and cell lines

Content and secretion of proANP and PAM were determined for atrial cultures after a number of drug treatments. For secretion, atrial cultures plated into a 96 well plate (on average, 1 atrium/well; confluent at the time of the experiment) were rinsed with 200 μl warm complete serum-free medium (CSFM) containing 100 μg/ml bovine serum albumin (Sigma A6003) (9), and incubated for 2h in 50 or 100 μl of the same medium. Medium was centrifuged at 80 x g for 5 min, and the supernatant saved with 0.3 mg/ml phenylmethylsulfonyl fluoride and protease inhibitor mix (9). Some drugs were administered only during the 2h secretion experiment: Brefeldin A (7.1 μM from a methanol stock; Sigma, B7651); Golgicide A (10 μM from a DMSO stock; Cayman, #18430), blebbistatin (10 μM from a DMSO stock; Sigma, B0560). Other drugs were administered into the growth medium for a pretreatment and then secretion was monitored into CSFM: cycloheximide (10 μM, 1h pretreatment); NH_4_Cl/methylamine (10 mM each, up to 12h pretreatment); MG-132 (5 and 10 μM, up to 12h pretreatment; Tocris, #1748); Bortezomib (32.5 nM, Selleckchem, PS-341); E-64 (20 μM; Tocris, #5208); PD150606 (50 μM; Tocris, #1269). Cell extracts were subjected to Western blot analysis for proANP, PAM and Gapdh; for denser cultures, cell extracts were also analyzed for cardiac troponin T; media were analyzed for proANP. Cell extracts and media were analyzed at the same time, allowing the calculation of secretion rates (% cell content secreted per h). The efficacy of Brefeldin A and Golgicide A was verified by monitoring the secretion of prohormone convertase 1 and proopiomelanocortin products by AtT-20 cells treated with the same dose of each drug for the same amount of time (18).

### Preparation and analysis of coated vesicles

A simplified protocol was developed from methods used to process larger amounts of tissue (19, 20). Atria from 10 to 12 adult mice (male and female; ∼100 mg wet weight) were rinsed in PBS, minced and placed into 1.0 ml of ice cold 0.1 M Na MES, 1 mM EGTA, 0.5 mM MgCl_2_, pH 6.5 (MES, pH6.5 buffer) containing protease inhibitor cocktail and dispersed using a Polytron. A low speed supernatant was saved after centrifugation at 4,000 xg for 20 min; the resulting pellet was re-extracted in 0.5 ml of the same buffer. The pooled low-speed supernatants were centrifuged at 8,000 xg for 20 min and the pellet discarded. Centrifugation of the pooled supernatants at 435,000 xg for 15 min yielded a high-speed pellet that contained coated vesicles and a supernatant that was discarded. The high-speed pellet was suspended in 0.4 ml MES, pH 6.5 buffer using a Dounce homogenizer and applied to a step gradient composed of MES, pH 6.5 buffer containing increasing concentrations (wt/vol) of sucrose: 60% (0.2 ml), 55% (0.2 ml), 50% (0.4 ml), 40% (0.4 ml), 30% (0.4 ml), 20% (0.4 ml). Fifteen 150 μl samples were collected from top to bottom. For Western blot analysis, an equal volume of each fraction was analyzed. For negative stain electron microscopy, a 5 μl aliquot of gradient fraction #8 was applied to a glow-discharged 400 mesh carbon coated copper grid (Electron Microscopy Sciences, Hatfield, PA); excess liquid was removed after 30 to 60 sec, the grid was rinsed with water, placed onto a drop of freshly made 1% uranyl acetate in water for 30 sec and air dried. Images were captured using a Hitachi H-7650 transmission electron microscope (Hitachi High Technologies Corporation, Tokyo, Japan) operating at 80 kV.

### Expression of proANP-Emerald and proNPY-GFP in HEK293 and PAM/HEK cells

HEK293 cells were obtained from Dr. Jeremy Nathans (Johns Hopkins University). HEK293 cells stably expressing rat PAM1 were generated by transfection (21) and are referred to as PAM/HEK cells. A vector encoding proANP-Emerald was obtained from Dr. Edwin Levitan (University of Pittsburgh). The vector encoding proNPY-GFP was constructed as described (18). HEK293 and PAM/HEK cells were transiently transfected using TransIT-2020 (Mirus Bio, Madison WI). For biochemical experiments, cells plated into plastic dishes were transfected 24 h later with 1.0 µg DNA/1.5 µl Transit 20:20 in Optimem (ThermoFisher). Secretion experiments were carried out 24 h after transfection, with media collected for 2h. For fluorescence microscopy, HEK and PAM/HEK cells were plated onto polylysine-coated 12 mm diameter, gelatin-coated German glass coverslips (#72293-01, Electron Microscopy Sciences, Hatfield, PA); cells were fixed for 20 min with 3.7% formalin in PBS. Following permeabilization with 0.1% TX-100 for 10 min, cells were blocked with bovine serum albumin and then incubated simultaneously with an antibody to PAM and an antibody to ERGIC-53 or to GM130 (**Supplemental Table 1**). For categorization of soluble cargo protein distribution as Diffuse, Peri-nuclear and Condensed, coverslips were systematically scanned and images of the fluorescently tagged cargo were acquired, along with images of Hoechst-stained nuclei. Coded images were analyzed by a blinded observed; between four and six groups of 50 to 100 cells were analyzed; the final data represent the average ± standard error of the four to six groups. Confocal images were acquired using a Zeiss LSM 880 confocal microscope with a 63X/1.4 NA Plan-Apochromat DIC M27 oil objective. Images were recorded sequentially with four separate detectors to avoid any cross talk among fluorochromes. Images were analyzed using Image J (https://imagej.nih.gov/ij/).

### Statistical analyses

Pairwise comparisons were performed using Student’s t-test (Excel or Open Office). ANOVAs were performed using GraphPad Prism 8. Linear best fit lines were assigned using Open Office.

## SUPPLEMENTARY MATERIAL

### Supplementary Figures

**Fig. S1.**
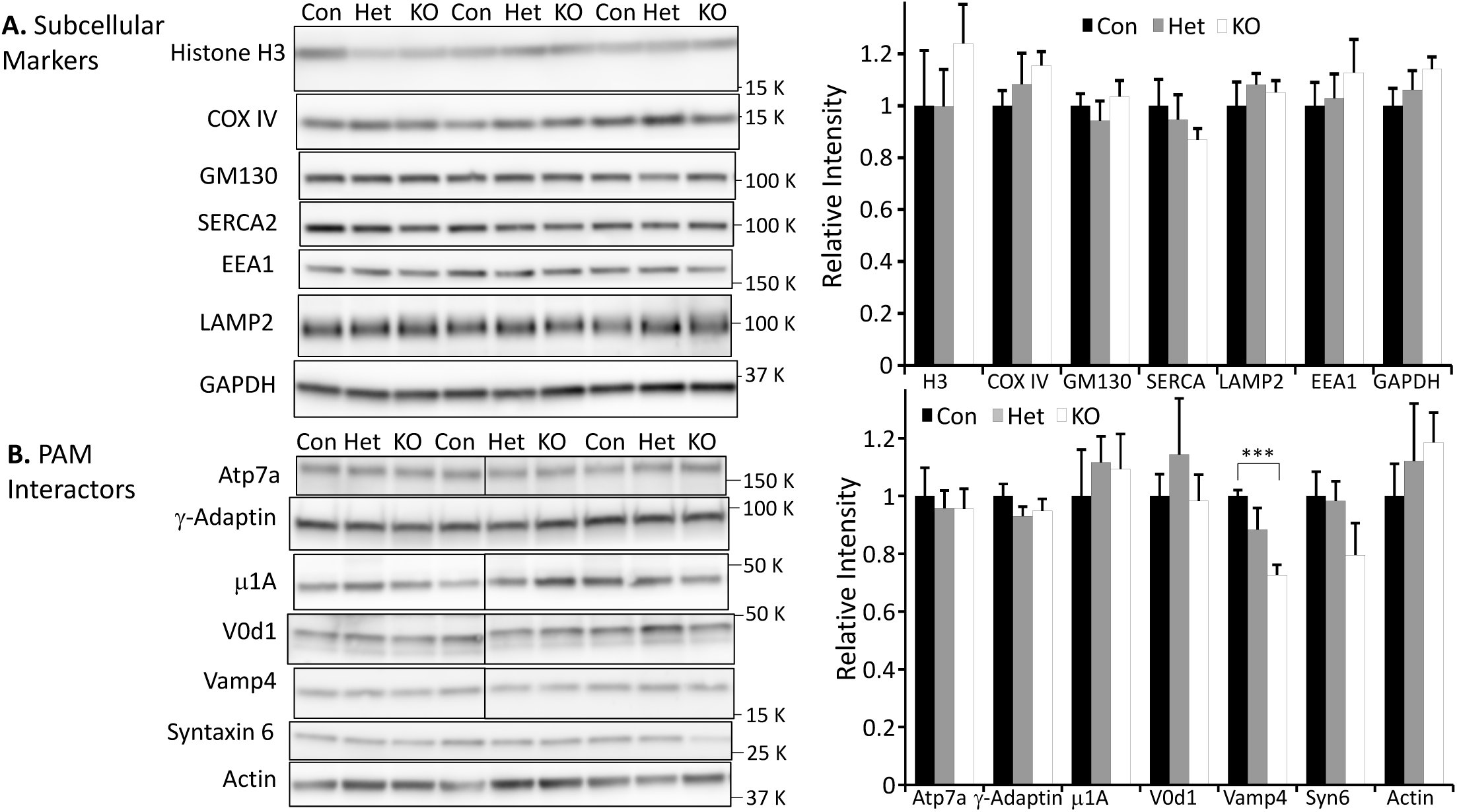
A. Levels of subcellular markers and PAM interactors. Expression of markers for nuclei (Histone H3), mitochondria (COX IV, cytochrome c oxidase subunit 4), Golgi (GM130, 130 kDa *cis*-Golgi matrix protein), sarcoplasmic/endoplasmic reticulum (SERCA, sarcoplasmic/endoplasmic reticulum calcium ATPase), early endosomes (EEA1, early endosomal antigen 1), lysosomes (LAMP2, lysosome associated membrane protein 2) and cytosol (GAPDH, glyceraldehyde 3-phosphate dehydrogenase) was evaluated in the total cell lysates analyzed in **Fig. 3A**. Average control intensities for each protein were normalized to 1.00 and Relative Intensities for Het and KO animals are shown. Antibody sources and concentrations are presented in **Supplementary Table 1**; n values were 5 or 6 for each genotype. Data for males and females did not differ and were pooled. Statistical analysis - error bars represent SEM; *, p<0.01; **, p<0.001, ***, p<0.0001. **B. PAM interactors.** The expression of proteins known to interact with PAM was evaluated in the same lysates. Based on studies in corticotrope tumor cells, HeLa and HEK293 cells, PAM interacts with the μ1A subunit of the adaptor protein 1 (AP1) complex, with Atp7a, a copper transporting ATPase, with the V0 subunit of the vacuolar proton pump (V-ATPase) and with actin (22-24). Levels of μ1A, γ-adaptin, Atp7a, V0d1 and actin were not affected by genotype. Trafficking proteins known to co-localize with PAM in corticotrope tumor cells include Vamp4 and Syntaxin 6 (25). Levels of Vamp4 were significantly decreased in the atria of *Pam*^Myh6-cKO/cKO^ mice. Vertical lines mark blots where intervening samples were deleted. ***, p-0.00003.

**Fig. S2.**
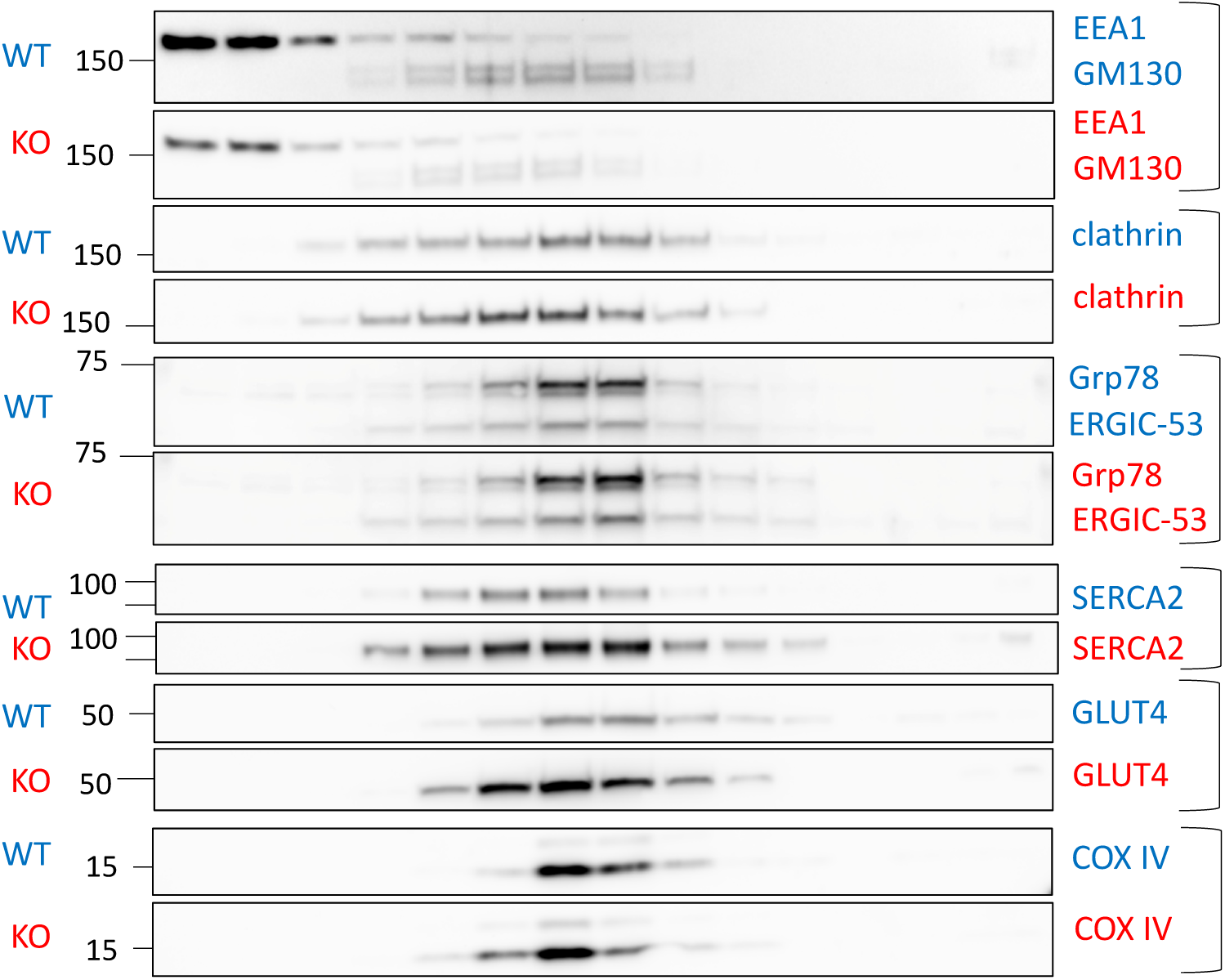
Marker protein localization in P3 gradients of control and *Pam*^Myh6-cKO/cKO^ atria. Gradients of P3 fractions prepared simultaneously from control and *Pam*^Myh6-cKO/cKO^ atria were analyzed for marker protein localization, as described for **Figs. 3B** and **3D**. No significant differences were observed in the localization of EEA1 (early endosome antigen 1), GM130 (a *cis*-Golgi matrix marker also known as Golga2), clathrin heavy chain (a marker for clathrin coated vesicles), Grp78 (78 kDa glucose regulated protein, an ER marker also known as BiP) and ERGIC-53 (also known as Lman1), SERCA2 (sarcoplasmic/endoplasmic reticulum calcium ATPase 2, an ER marker), GLUT4 (Glucose transporter type 4, insulin responsive, an endosomal/plasma membrane marker) and COXIV (Cytochrome c oxidase subunit 4, a mitochondrial marker).

**Fig. S3.**
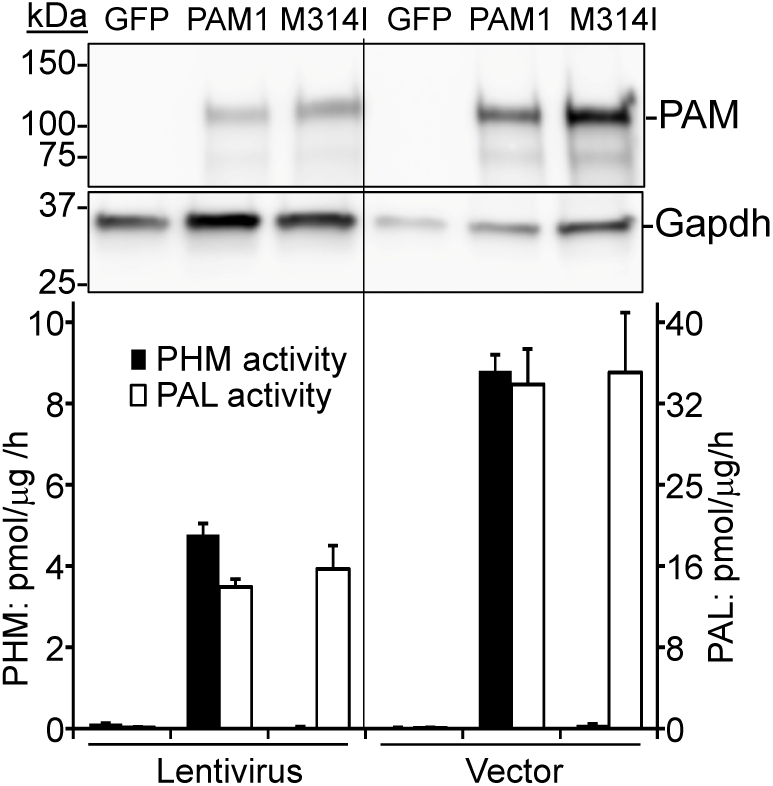
Validation of lentiviruses encoding PAM1 and PAM1/M314I. Densely plated pEAK-Rapid cells, a derivative of HEK293 cells, were transduced with lentivirus encoding GFP, PAM1 or PAM1 with a monooxygenase inactivating mutation (Met314Ile) (M314I) or transfected with expression vectors encoding the same proteins using TransIT-2020 (Mirus Bio, Madison WI) (26). **Upper Panels:** For enzyme assays, cells were extracted into 20 mM Na TES, pH 7.4 containing 100 mM mannitol and 1% TX-100. In order to verify the presence of intact PAM protein, extracts were analyzed by Western blotting using antibodies to PAM (JH629) and Gapdh. **Lower Panel:** For PHM and PAL assays, lysates were assayed with [^125^I]-Ac-Tyr-Val-Gly or [^125^I]-Ac-Tyr-Val-α-hydroxy-Gly and 0.5 μM of the corresponding unlabeled peptide, respectively (27). The Met314Ile mutation eliminated PHM activity without affecting PAL activity.

**Fig. S4.**
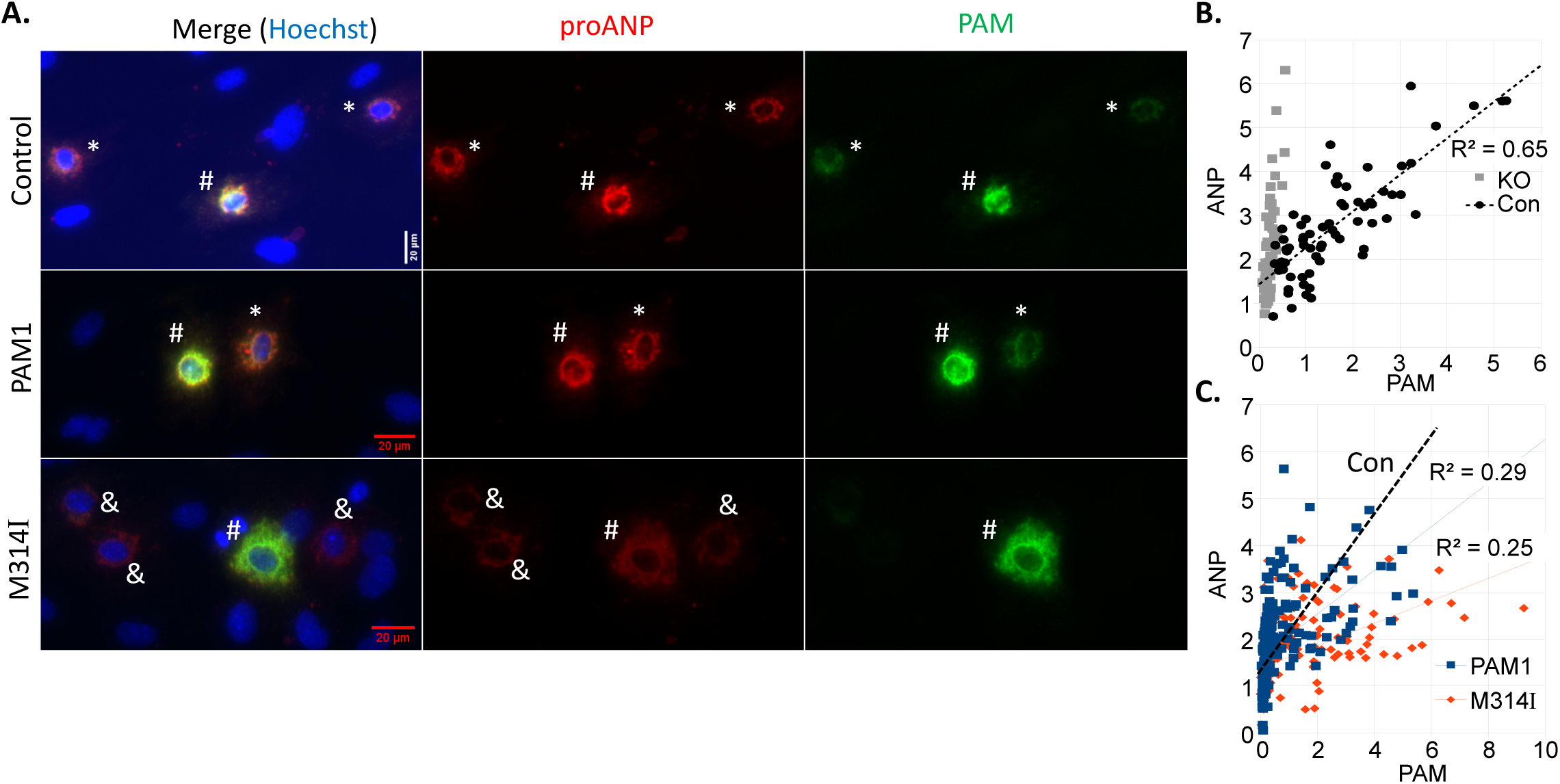
Rescue analysis using mouse monoclonal antibody to PAM and rabbit antibody to proANP. **A.** Atrial cultures (Control and *Pam*^Myh6-cKO/cKO^ (KO)) were prepared as described for **Fig. 4.** KO cultures were spinoculated with lentivirus encoding PAM1 or PAM1/Met314Ile at the time of plating. After 7 days *in vitro*, cultures were fixed and stained for PAM using a mouse monoclonal antibody (6E6 and Alexa Fluor 488 goat anti-mouse IgG) and for proANP using a rabbit antiserum directed to the N-terminal region of proANP ((8) and Alexa Fluor 633 goat anti-rabbit IgG); nuclei were visualized using Hoechst stain. KO cultures that had not been spinoculated exhibited no PAM staining and images are not shown. Atrial myocytes expressing higher (#) and lower (*) levels of PAM are marked; myocytes with ANP but no PAM are marked by a &. Staining intensities for individual cells were quantified after subtracting a locally determined background and data for Control and KO cultures (**B.**) and for PAM1 and M314I spinoculated cultures (**C.**) are plotted as described for **Fig.4.**

**Fig. S5.**
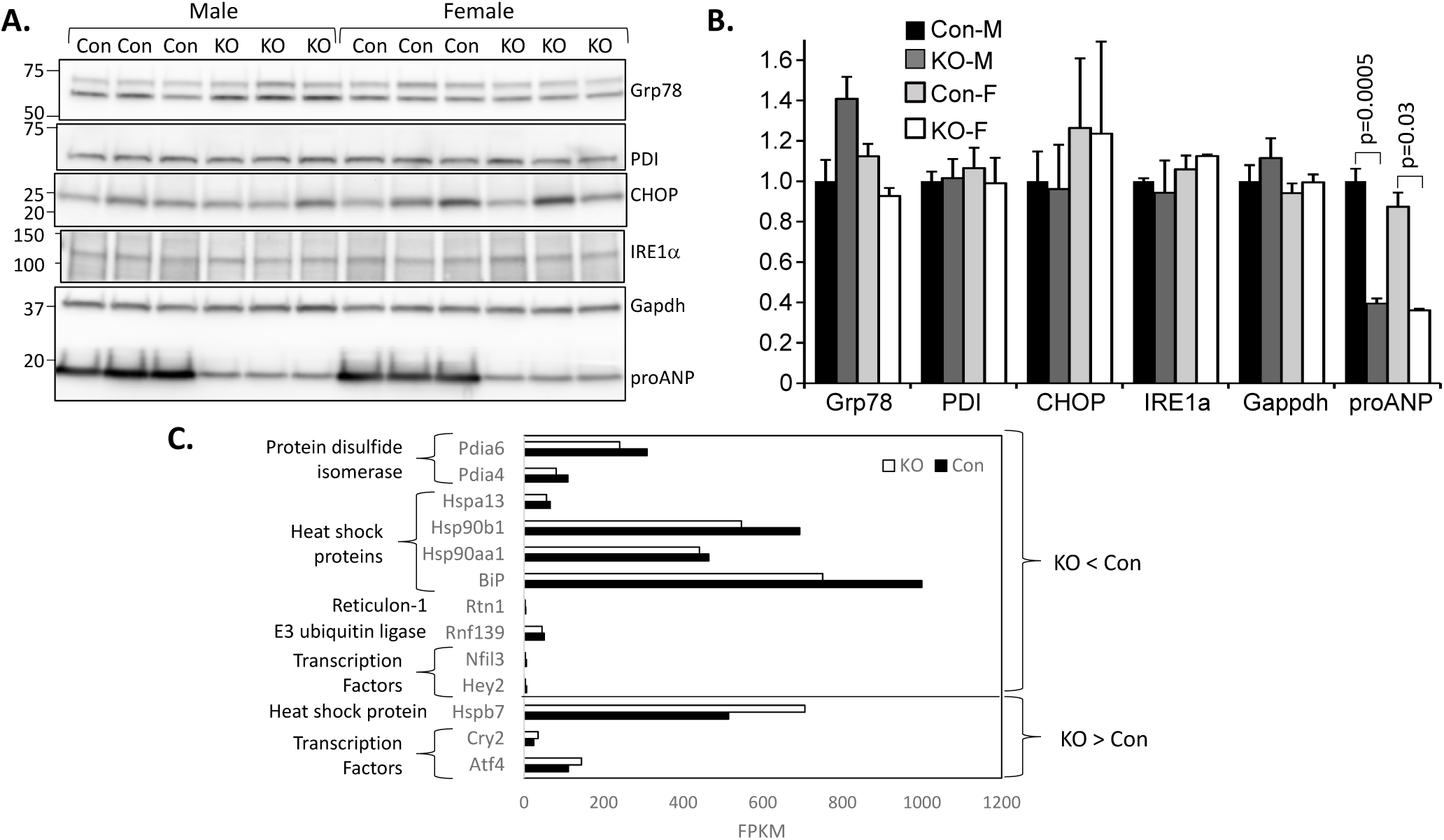
The absence of PAM does not trigger ER stress. **A.** The expression of proteins whose levels are known to rise as part of the ER stress response in cardiomyocytes (28-30) was evaluated using SDS-lysates prepared from control and *Pam*^Myh6-cKO/cKO^ adult male and female mice (n = 3/group). **B.** Data for each protein were normalized to the average level in control male mice. Levels of Grp78 (BiP/Hspa5), protein disulfide isomerase (PDI), CHOP (Ddit3) and inositol-requiring protein 1 (IRE1α/Ern1) did not differ significantly. ProANP levels dropped as expected, with no change in Gapdh levels. **C.** A set of 56 transcripts whose expression has been shown to vary in response to ER stress in cardiomyocytes (28, 29), β-cells or provasopressin-producing neurons (31-35) was compiled (**Supplementary Table 2**). Transcripts whose levels differed significantly (DESeq2 p-value < 0.05) in wildtype and *Pam*^Myh6-cKO/cKO^ atria (1) are shown, with average wildtype and KO FPKM values. Included in the list were ER localized heat shock proteins, protein disulfide isomerases, E3 ubiquitin ligases and reticulon proteins known to play a role in ER-phagy (31, 36). The decreased levels of multiple heat shock proteins, an ER localized E3 ubiquitin ligase (Rnf139), two protein disulfide isomerases and reticulon 1 do not suggest that lack of PAM caused generalized ER stress or induced ER-phagy (33). Four of the down-regulated transcripts are part of a proteostasis network that includes ER resident (Hspa5, Pdia4 and Pdia6) and cytosolic (Hsp90aa1) components interconnected by calnexin (Canx), a calcium-binding ER resident protein whose expression was also reduced in the *Pam*^Myh6-cKO/cKO^ atrium (503 FPKM in wildtype vs. 422 FPKM in *Pam*^Myh6-cKO/cKO^, p=0.0095). Without the need to assist in the folding of a major secretory pathway protein (PAM), expression of relevant chaperones and disulfide isomerases was reduced.

**Fig. S6.**
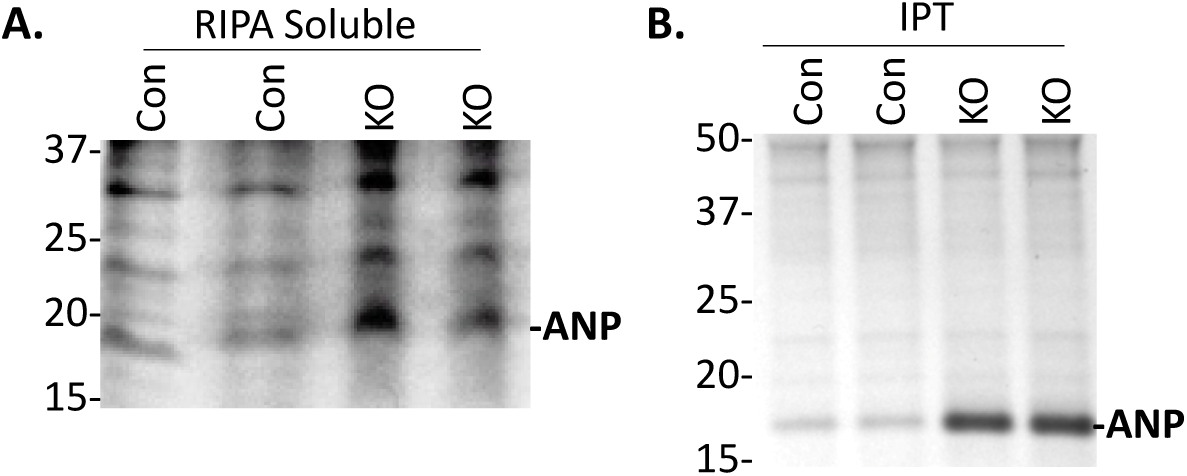
Metabolic labeling of proANP in Control (wildtype) and *Pam*^Myh6-cKO/cKO^ (KO) atria. Fragments of adult atrial tissue were incubated in methionine-free medium containing [^35^S]-Met and [^35^S]-Cys as described in Methods. **A.** RIPA extracts were analyzed without fractionation; its prevalence allows quantification of proANP without further purification (**Fig. 3** and (37)). **B.** After pre-adsorption with Protein A, a rabbit polyclonal antibody to the N-terminal region of proANP was used to isolate proANP. Quantitative data for total lysates and proANP immunoprecipitates are shown in **Fig. 5A**.

**Fig. S7.**
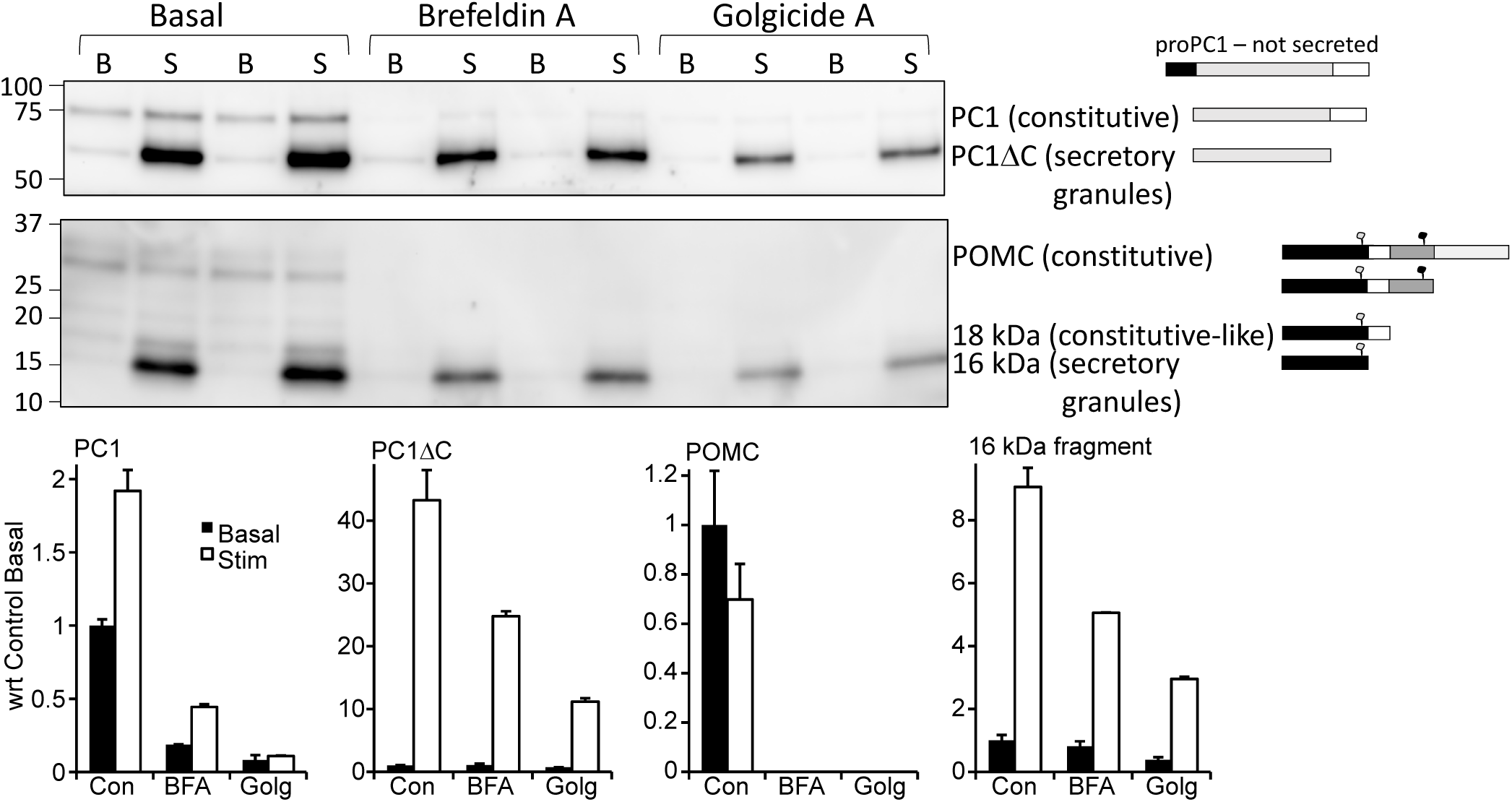
Validation of BFA and Golgicide A efficacy using AtT-20 cells. Duplicate wells of AtT-20 cells were rinsed in CSFM and incubated in CSFM or in CSFM containing 7.1 μM BFA or 10 μM Golgicide A for three sequential 30 min periods; spent media were harvested after each 30 min incubation and cells were harvested after the third incubation. During the final 30 min incubation, the media also contained 2 mM BaCl_2_, a potent stimulator of the regulated secretory pathway in AtT-20 cells (18). Equal aliquots of the spent media from period 1 (Basal, B) and period 3 (Stimulated, S) were fractionated by SDS-PAGE. Antisera to prohormone convertase 1 (JH888) and POMC (Georgie: the 16kDa fragment region) were used to compare products secreted basally and in response to secretagogue (BaCl_2_); diagrams to the right of the blots identify the major products generated from each precursor. Intact proPC1 is not secreted; PC1ΔC is generated by removal of the C-terminal region, which activates the enzyme (38). PC1ΔC is stored in secretory granules; very little is released in the absence of secretagogue. A substantial amount of PC1 is basally secreted. As expected, both BFA and Golgicide A had a more profound effect on the basal secretion of PC1 than on the stimulated secretion of PC1ΔC. Intact POMC (the two bands differ by the presence and absence of an N-linked oligosaccharide in the C-terminal region of ACTH) is released in the absence of secretagogue and is not responsive to secretagogue addition. Both BFA and Golgicide A almost eliminated the basal secretion of POMC. The 16 kDa fragment produced from POMC is stored in secretory granules; only small amounts are secreted basally, but BaCl_2_ produced a more than 8-fold increase in its secretion. While BFA and Golgicide A each had an inhibitory effect on the secretion of 16 kDa fragment, the effect was modest compared to their almost total inhibition of POMC release.

**Fig. S8.**
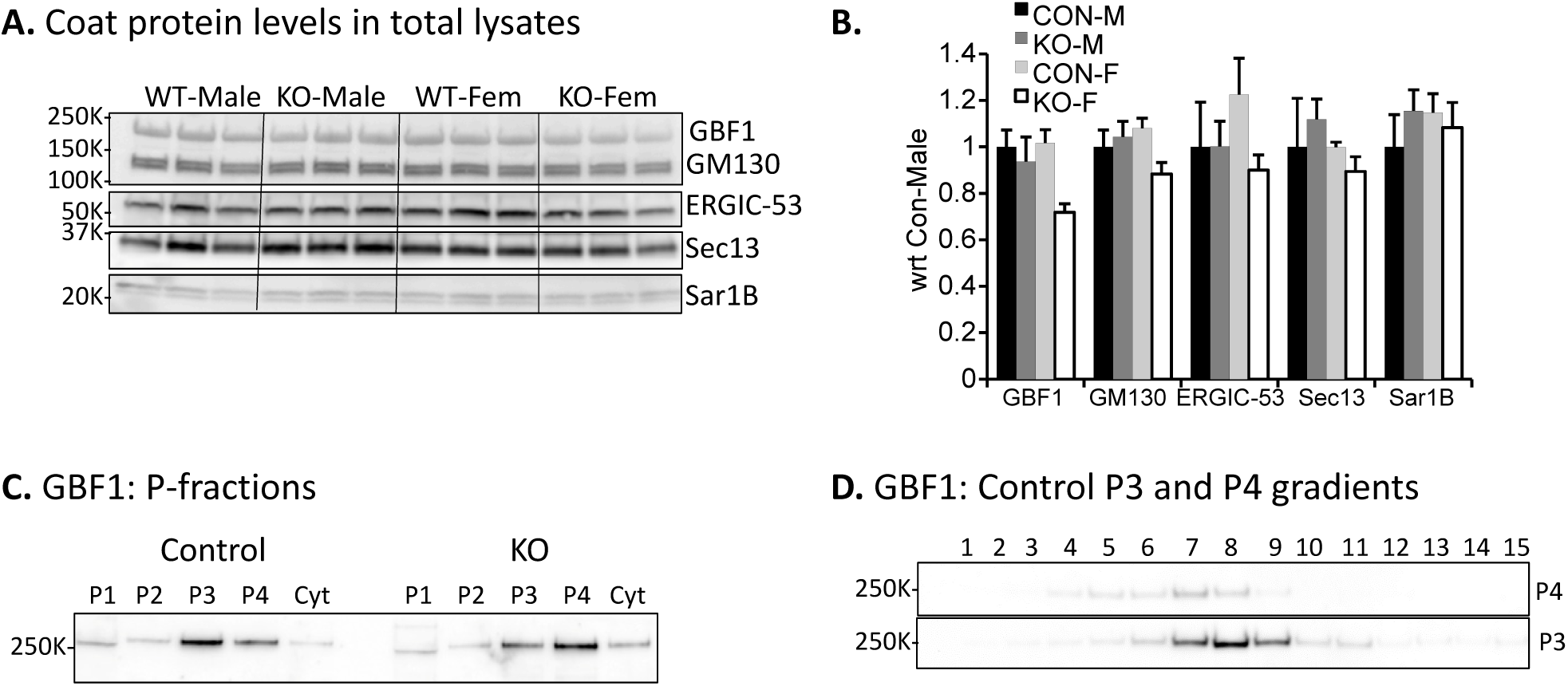
Coated vesicle/trafficking protein levels in Control and KO Atria. **A.** Protein levels in lysates prepared from Control and KO adult atria were evaluated by Western blot analysis as described for **Fig. S5. B.** Levels of GBF1, Sec13, Sar1B, GM130 and ERGIC-53 did not differ by genotype or by sex; data were normalized to average values for Control males. **B.** As described for **Fig. 3C**, subcellular fractions prepared from Control and KO atria were analyzed for their content of GBF1. **C, D.** As described for **Fig. 3C, D**, P3 and P4 fractions prepared from Control atria were subjected to sucrose gradient fractionation; GBF1 was localized to fractions enriched in ER/Golgi markers.

**Fig. S9.**
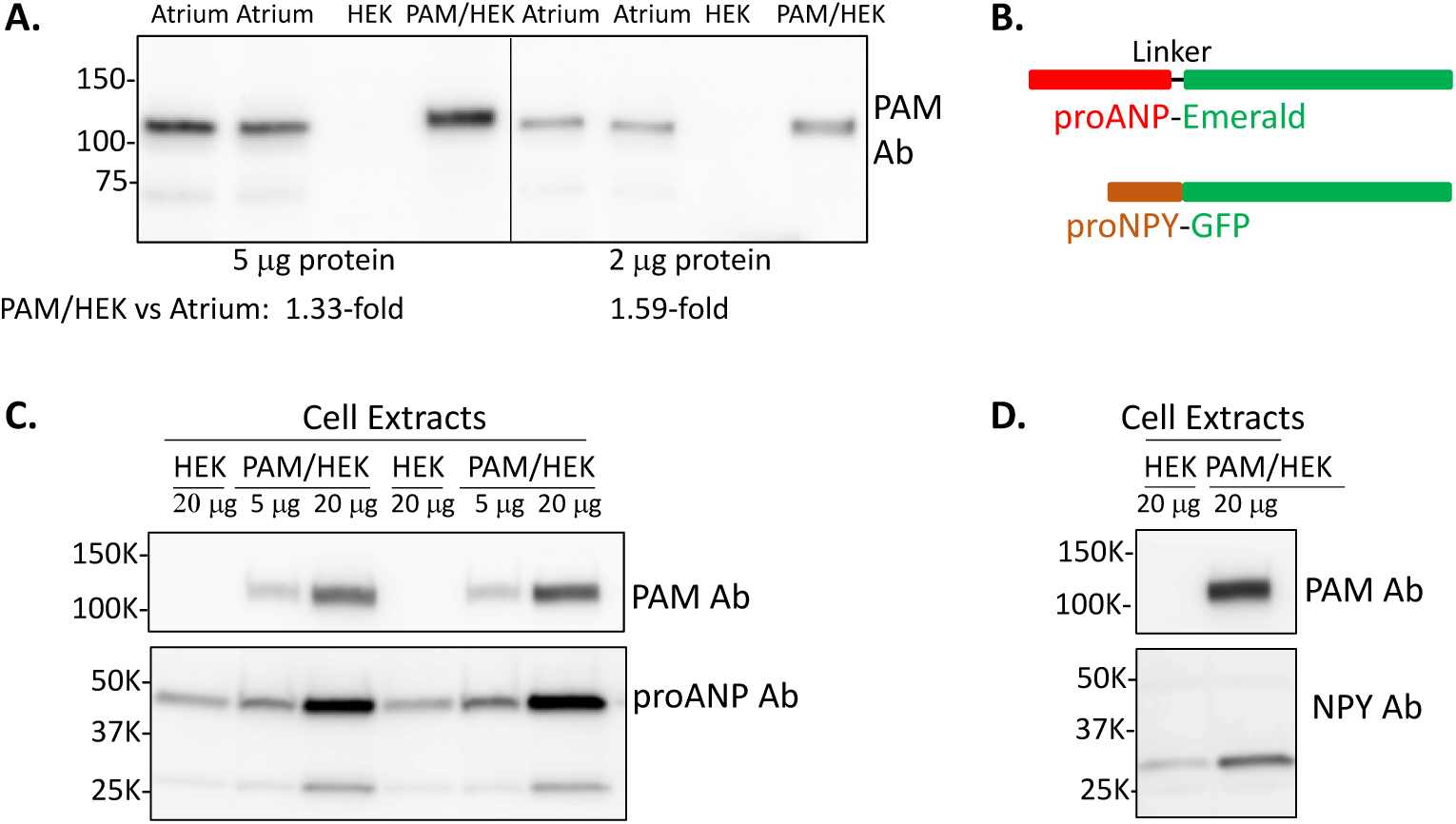
PAM levels in atrium vs. PAM/HEK cells and transient expression of proANP-Emerald and proNPY-GFP in HEK and PAM1/HEK cells. **A.** SDS lysates prepared from adult mouse atria (n = 2), HEK cells and PAM/HEK cells were analyzed for PAM content; the panel to the left shows data obtained using 5 μg of protein and the panel to the right shows data obtained using 2 μg protein. PAM levels in PAM/HEK cells were 1.33 and 1.59-fold higher than atrial levels. **B.** Diagrams show the structures of the mature proteins, proANP-Emerald and proNPY-GFP, produced following expression of vectors encoding preproANP-Emerald and preproNPY-GFP, respectively. HEK and PAM/HEK cells were harvested 24 h after transient transfection with vectors encoding preproANP-Emerald (**C**) or preproNPY-GFP (**D**). The regions of the membrane containing proteins above 75 kDa in apparent molecular mass were exposed to a monoclonal antibody specific for the C-terminus of PAM1 (**C** and **D**); rat PAM1, which is stably expressed in PAM/HEK cells was visible, but the endogenous levels of human PAM in HEK cells fall below the limit of detection using this antibody. The regions of the membrane containing proteins below 75 kDa in apparent molecular mass were exposed to proANP antibody (**C**) or to NPY antibody (**D**). While most of the proANP remained intact, a lower molecular weight product was detected in both HEK and PAM/HEK cells.

### Supplementary Tables

**Table S1.** Antibodies Used.

| Antigen (gene) | Antibody | Source | Reference |
| --- | --- | --- | --- |
| Actin | GTX14128 | GeneTex |  |
| γ-adaptin (Ap1g1) | 610385 | BD Transduction Labs | |
| ANP (Nppa) | proANP(1-16) | Dr. Chris Glembotski | (8) |
| ANP (Nppa) | ab190001 | Abcam |  |
| Atp7a/Mnk (Atp7a) | CT77 | AB_2827618 | (39) |
| BNP (Nppb) | bs-2207R-TR | Bioss |  |
| Carboxypeptidase E (Cpe) | C-terminal, Ab139 | Dr. Lloyd Fricker | (40) |
| CHOP (Ddit3) | L63F7 | Cell Signaling Technologies |  |
| Clathrin, H chain (Cltc) | 610499 | BD Transduction Labs |  |
| α-COP (Copa) | Ab181224 | Abcam | |
| COXIV (Cox4i1) | 11242-1 | Proteintech |  |
| EEA1 (Eea1) | C45B10 | Cell Signaling Technologies |  |
| ERGIC-53 (Lman1) | 13364-1-AP | Proteintech |  |
| GAPDH (Gapdh) | MAB374 | EMD Millipore Corp |  |
| GBF1 (Gbf1) | ab189512 | Abcam |  |
| GLUT4 (Slc2a4) | 66846-1 | Proteintech |  |
| GM130 (Golga2) | 610822 | BD Transduction Labs |  |
| Grp78/BiP/HSA5 (Hspa5) | PA5-29705 | Invitrogen |  |
| Histone, H3 (H3.1-291) | 9715 | Cell Signaling Technologies |  |
| IRE1-α (Ern1) | 14C10 | Cell Signaling Technologies | |
| LAMP2 (Lamp2) | ab13524 | Abcam |  |
| μ1A (Ap1m1) | 12112-1-AP | Proteintech | |
| PAM (Pam) | JH629 | AB_2721274 | (41) |
| PAM (Pam) | mAb 6E6 | AB_28277619 | (42) |
| PDI | C81H6 | Cell Signaling Technologies |  |
| Prohormone convertase 1 (Pcsk1) | JH888 | AB_2802129 | (43) |
| Proopiomelanocortin (Pomc) | 16K fragment | AB_2721275 | (44) |
| Sar1B (Sar1b) | Ab155278 | Abcam |  |
| Sec13 (Sec13) | 15397-1-AP | Proteintech |  |
| SERCA2 (Atp2a2) | ab219173 | Abcam |  |
| Syntaxin6 (Stx6) | 610635 | BD Transduction Labs |  |
| Troponin T (TnTc) (Tnnt2) (cardiac) | 15513-1-AP | Proteintech |  |
| V0d1 (Atp6v0d1) | 18274-1-AP | Proteintech |  |
| VAMP4 (Vamp4) | 136 002 | Synaptic Systems, Gottingen |  |

***Table S2. RNASeq data for genes of interest.*** *Excel file.*

FPKM values and DESeq2 data from Powers et al (2019) PNAS 116:20169. Transcripts known to be part of the ER Stress Response Pathway (Page 1), Proteasome Function (Page 2), the Secretory Pathway (Page 3) and Antibodies used in Western Analyses (Page 4) are compiled.

